# WWP2 MEDIATES THE METABOLIC REPROGRAMMING OF RENAL MYOFIBROBLASTS TO PROMOTE KIDNEY FIBROSIS

**DOI:** 10.1101/2023.08.22.554242

**Authors:** Huimei Chen, Ran You, Jing Guo, Wei Zhou, Gabriel Chew, Nithya Devapragash, Jui Zhi Loh, Loreto Gesualdo, Yanwei Li, Yuteng Jiang, Elisabeth Li Sa Tan, Shuang Chen, Paola Pontrelli, Francesco Pesce, Jacques Behmoaras, Aihua Zhang, Enrico Petretto

## Abstract

Renal fibrosis is a common pathological endpoint in chronic kidney disease (CKD) that is challenging to reverse. Although myofibroblasts are mainly responsible for the accumulation of a fibrillar collagen-rich extracellular matrix (ECM) in fibrotic kidney, recent studies have unveiled their diversity in terms of proliferative and fibrotic characteristics. This diversity could be linked with the existence of different metabolic states, and myofibroblast metabolic reprogramming may contribute to the pathogenesis and progression of renal fibrosis. Here, we reveal an unexpected role of the E3 ubiquitin-protein ligase WWP2 in the metabolic reprogramming of myofibroblasts during renal fibrosis. The tubulointerstitial expression of WWP2 contributes to the progression of fibrosis in CKD patients, and in pre-clinical murine models of CKD. WWP2 deficiency increases fatty acid oxidation and activates the pentose phosphate pathway, boosting mitochondrial respiration at the expense of glycolysis. This concurrently promotes myofibroblast proliferation and halts pro-fibrotic activation, reducing the severity of kidney fibrosis. Mechanistically, WWP2 suppresses the transcription of PGC-1α, a metabolic mediator shaping myofibroblast fibrotic response. Pharmacological interventions targeting PGC-1α reverse the effects of WWP2 on fibrotic myofibroblasts. These findings demonstrate the influence of WWP2 on essential metabolic pathways involved in fibrogenesis, uncovering the WWP2-PGC-1α axis that orchestrates the metabolic reprogramming of myofibroblasts during renal fibrosis. Our study presents a potential novel target for therapeutic intervention in the treatment of chronic kidney disease.

**Graphical abstract:** 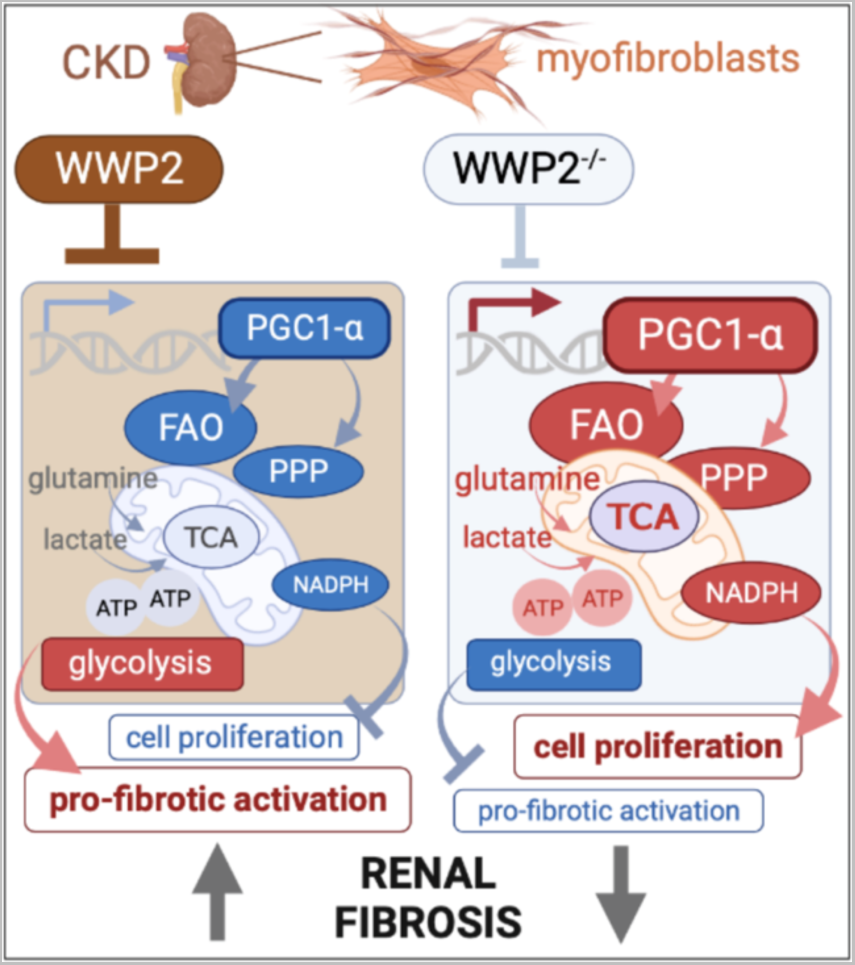

**Highlights:**

- WWP2 expression is elevated in the tubulointerstitium of fibrotic kidneys and contributes to CKD pathogenesis and progression.
- WWP2 uncouples the pro-fibrotic activation and cell proliferation in renal myofibroblasts.
- WWP2 controls mitochondrial respiration in renal myofibroblasts through the metabolic regulator PGC-1α
- Myofibroblast metabolic reprogramming mediates the effect of WWP2 on fibrotic myofibroblasts.

## Introduction

Chronic kidney disease (CKD) is a global health concern and has become one of the leading causes of mortality worldwide. CKD-associated deaths have surged over the past two decades [1]. Renal fibrosis is a common pathological pathway in CKD irrespective of the underlying aetiology [2]. It is broadly characterized by the accumulation of a fibrillar collagen-rich extracellular matrix (ECM) in the tubulointerstitial space, which eventually results in end-stage renal disease [3]. During renal fibrosis, interstitial fibroblasts transdifferentiate into myofibroblasts, which contribute to ECM accumulation. Recent studies highlights the complexity of this phenomenon as the fibrotic (or inflamed) kidney contains a variety of pro-fibrotic cells [4, 5], suggesting extensive fibroblast heterogeneity [6]. Although alpha- smooth-muscle actin (α-SMA) is commonly used as a specific marker for activated myofibroblasts[7, 8], only around 50% of α-SMA^+^ cells express Col1a1 (collagen type I alpha 1), and the other half contribute minimally to ECM deposition during renal fibrogenesis [9]. Cell proliferation is another key response to renal injury [10], and it is observed along the differentiation trajectories of myofibroblast activation [11]. This suggests a tight association between the proliferative state of a fibroblast and its subsequent activation and ECM production [12]. However, other mechanisms independent of cell activation have been reported for myofibroblast proliferation [13, 14]. Decoding the complex interplay and mechanisms underlying myofibroblasts activation, proliferation, and ECM production is crucial for understanding the development of pathological renal fibrosis in CKD [7].

Fibrometabolism is an emerging avenue of research [15], and metabolic reprogramming is increasingly considered as a fundamental process for myofibroblast proliferation and activation in different organs where tissue fibrosis occurs [6, 16]. During skin wound healing, disruptions of the enzymes involved in glycolysis and the fatty acid oxidation (FAO) display their reciprocal effects on ECM upregulation and downregulation, respectively [17]. In lung fibrosis, the increase in aerobic glycolysis represents a critical factor for myofibroblast activation, particularly when coupled with profibrotic TGF-β signalling activation [18, 19]. *In vitro* studies showed how suppression of myofibroblast proliferation is linked to a metabolic flux rerouting from the pentose phosphate pathway (PPP) toward glycolysis, coupled with an increased expression of ECM proteins [20]. Elevated reactive oxygen species (ROS) is also reported to harm cardiac myofibroblasts [21], while a reduction in the generation of NADPH is observed in pulmonary myofibroblasts [22]. Considerable controversy remains regarding the heterogeneity of myofibroblasts and complexity of metabolism; however, these studies associate myofibroblast metabolism reprogramming with their proliferation and ECM synthesis in relation to fibrogenesis, suggesting potential new approaches for antifibrotic therapy.

Focusing in particular on kidneys, altered energy metabolic activities have been observed in injured tissue and associated with fibrotic progression in both CKD patients and murine models of CKD [23]. Moreover, kidney function can be improved by enhancing FAO and oxidative phosphorylation (OXPHOS) [24], while ramping down glycolysis can effectively slow down the progression of kidney fibrosis [25]. The potential participation of mitochondrial function and dynamics has been proposed in renal myofibroblast activation [26]. During kidney fibrosis, dysregulated myofibroblast metabolism may be critical for shaping cellular proliferation and pro-fibrotic response [27]. Beyond the setting of myofibroblasts in kidney, there is an increasing emphasis on the metabolism programming in proximal tubule (PT) cells [28–30], which is partly attributable to PT cells containing one of the highest mitochondria density in the body. Yet, the underlying regulators of metabolism within the kidney, particularly concerning myofibroblasts remain poorly defined. This lack of clarity might be a result of the complexity of the kidney anatomy and the heterogeneity of myofibroblasts in comparison to other tissues.

The peroxisome proliferator-activated receptor gamma coactivator 1-alpha (PGC-1α) has emerged as a common regulator of the metabolic processes underlying fibrosis in multiple tissues [31]. The regulation of myofibroblast metabolism by PGC-1α has been shown in wound healing, lung fibrosis, and cardiac remodeling, where it affects mitochondrial biogenesis, oxidative phosphorylation, and secretory state of myofibroblasts [32–34]. In the kidneys, low PGC-1α activity has been observed in association with CKD, and approaches to increase PGC-1α activity may pave the way for developing novel nephroprotective therapies [35]. PGC-1α is known to contribute to fibrotic progression in multiple models of kidney disease via its action outside the stromal compartment [35–37]. However, numerous studies detail the function of PGC-1α in PT cells in renal fibrosis, but its impact on renal myofibroblasts remains relatively unexplored. PGC-1α is reported to maintain basal mitochondrial function of renal PT cells and other subcellular processes such as mitochondrial biogenesis, dynamics, and mitophagy [38, 39]. PGC-1α also regulates PT metabolism processes including FAO and OXPHOS, that fine-tune kidney fibrosis progression [28, 29].

Experimental evidence supporting the myofibroblast metabolic reprogramming in the setting of renal fibrosis is only just beginning to emerge [6]. Here, we focus on myofibroblasts, shed light on their heterogeneity and assessed their metabolic regulation in fibrotic kidneys. We also postulate that metabolic reprogramming plays an important role in the fibrotic response of myofibroblasts during CKD. We previously identified the HECT domain E3 ubiquitin-protein ligase WWP2 as a regulator of myofibroblast pro-fibrotic activation in cardiac fibrosis [40]. Giving the wide-spread tissue expression and master regulatory role of WWP2 in orchestrating pro-fibrotic ECM network [40, 41], we hypothesised that WWP2 regulates myofibroblast phenotypes and tissue fibrosis also in kidney disease. In this work we first identify WWP2 up-regulation in tubulointerstitium of kidney from human CKD patients and pre-clinical mouse models of CKD. Integrating single-cell transcriptomics with metabolic assay, we then elucidate the role of WWP2 in driving myofibroblast metabolic reprogramming in kidney fibrosis, with a specific focus on its mechanism targeting PGC-1α transcription. We further propose that this WWP2-PGC-1α-axis in myofibroblasts regulates mitochondrial energy metabolism predominantly through FAO process, which underlies the change in renal myofibroblasts proliferation and ECM production, and, ultimately, renal fibrosis. These data identify WWP2 as key regulator of the metabolic reprogramming of myofibroblasts, and suggest that WWP2 contributes to maintain the balance between myofibroblast proliferation and activation, thereby playing a pivotal role in renal fibrosis during CKD.

## Results

### Tubulointerstitial expression of WWP2 is associated with severity of renal fibrosis

We investigated the expression of WWP2 in kidney biopsies obtained from patients with CKD and show that WWP2 was mainly expressed within the tubulointerstitial space, co-localized with fibrotic lesions (Supplementary Figure 1A). WWP2 expression was more pronounced in kidneys with severe fibrosis compared to those with mild fibrosis (Figure 1A). In a multi-centre cohort of CKD patients (n=133, See Supplementary Table 1), we found a significant correlation between the interstitial WWP2-positive area and fibrosis assessed by Sirius Red staining in the tubulointerstitial region (*r*=0.274, P=0.001, Figure 1B), which was further confirmed in two cohorts of patients with IgA nephropathy (IgAN) (Supplementary Figure 1B). Blood urea nitrogen (BUN) and serum creatinine levels were recorded in a subset of patients, and used to evaluate kidney function [42]. We detected a positive correlation between WWP2 expression in the tubulointerstitial region and the impairment of renal function (Supplementary Figure 1C), but not with proteinuria (Supplementary Figure 1D). Furthermore, patients with moderate CKD (stage 3-4) showed higher expression of interstitial WWP2 when compared with those having mild CKD (stage 1-2), regardless of the original insult leading to the kidney damage (Figure 1E). There was no significant association between the glomerular WWP2 expression and the extent of glomerular fibrosis [43], kidney function, and proteinuria in CKD patients (Supplementary Figure 1E-H).

**Figure 1.**
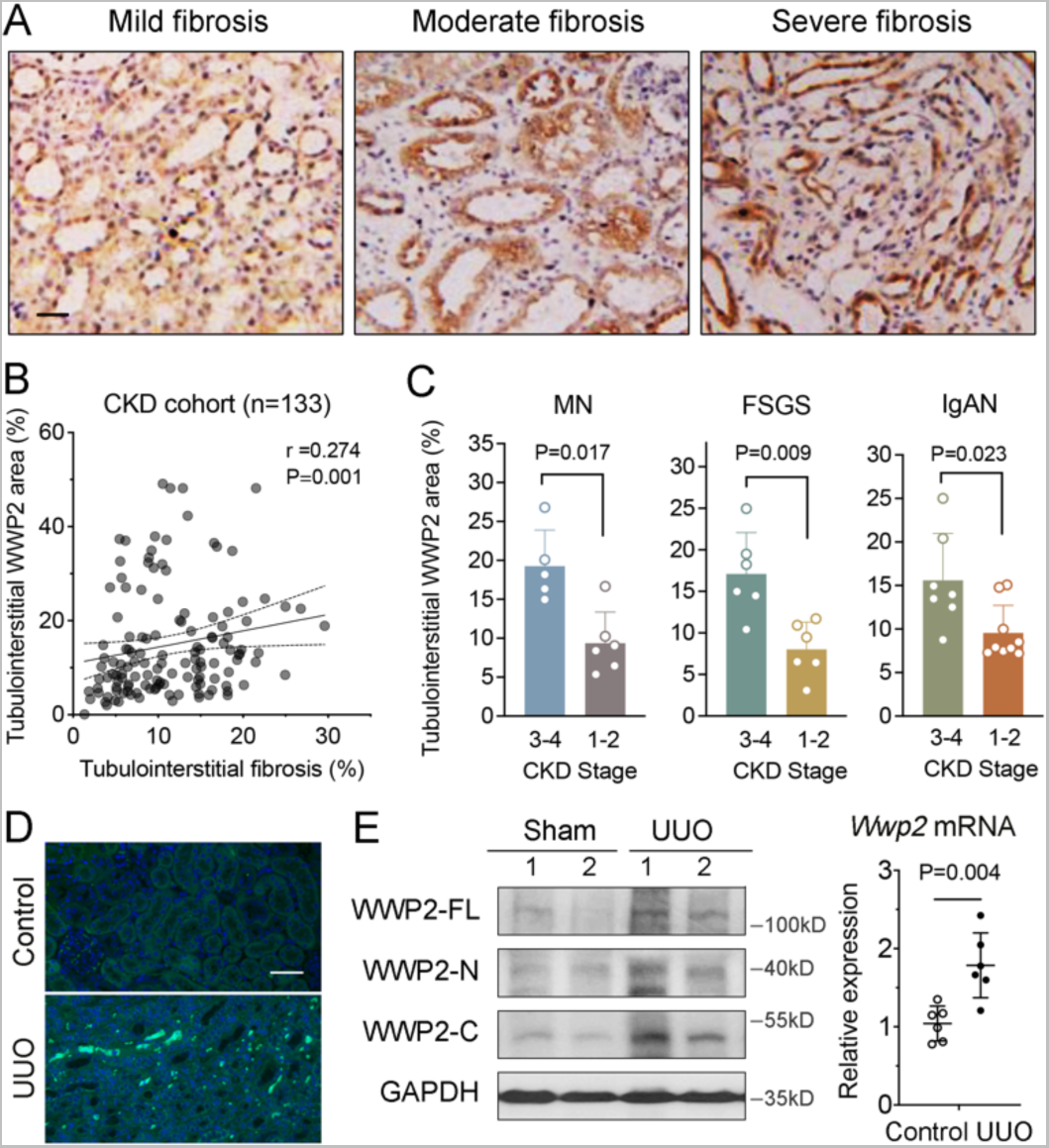
The tubulointerstitial WWP2 expression is positively associated with renal fibrosis in chronic kidney disease (CKD) patients and UUO-mice. (A) Representative immunostaining images of WWP2 in tubulointerstitial area from human renal biopsy samples, illustrating mild, moderate and severe interstitial fibrosis, respectively (n=133 patients with CKD recorded, see Supplementary Table 1). Scale bars, 30 μm. (B) Positive correlation between tubulointerstitial WWP2-positive area and fibrosis-positive area, as determined by immunostaining and Sirius Red staining (see Supplementary Figure 1A) with ImageScope. Each data point represents a measurement of an individual section obtained from each patient with CKD (n=133). (C) The WWP2-positive tubulointerstitial area (percentage) in renal biopsy samples increases as CKD progresses from stage 1-2 to stage 3-4, including renal biopsy samples from patients with membranous nephropathy (MN) (n=11, *P*=0.017), focal segmental glomerulosclerosis (FSGS)(n=12, *P*=0.009) and IgA nephropathy (IgAN) (n=16, *P*=0.023). P-values calculated by two-tailed Mann-Whitney U test. (D) Representative immunostaining images of WWP2 in tubulointerstitial kidneys from UUO and control mice (n=5-10 images recorded for each mouse kidney). Scale bars, 50 μm. (E) The expression of WWP2 in kidney tissue from UUO and control mice. *Left*, representative western blotting for protein levels; *right*, mRNA expression changes were determined by RT-qPCR. n=6, each group, and values are reported and mean ± SD.

The unilateral ureteral obstruction (UUO) model is a commonly used pre-clinical murine model for studying tubulointerstitial fibrosis [44]. In this model, the intensity of WWP2 staining was higher in UUO compared to control kidneys (Figure 1D). This was supported by the relatively increased protein and transcript expression levels of WWP2 in fibrotic kidneys (Figure 1E). We generated a transgenic mouse line engineered to overexpress WWP2 (WWP2^Tg^) (Supplementary Figure 2A). Compared to controls (Control^Tg^), WWP2^Tg^ mice exhibited higher percentage of fibrosis in the kidney sections (Supplementary Figure 2B), presenting significantly more severe fibrotic lesions visualised by Masson’s trichrome staining. WWP2 overexpression also increased fibrotic ECM marker genes in UUO kidneys, as shown by the upregulation of Acta2, Co1a3 and Fn-EDA (Supplementary Figure 2C).

Collectively, the data in human fibrotic kidneys and pre-clinical fibrotic mouse model suggest an association between the tubulointerstitial WWP2 upregulation and fibrotic progression in CKD.

### WWP2 deficiency protects from renal fibrosis *in vivo*

We previously demonstrated that WWP2 mutant mice (WWP2^−/−^, carrying a 4-bp deletion in exon 2 that leads to disruption of WWP2-FL and WWP2-N isoforms), are protected from heart fibrosis [40]. Here, we confirm the deletion of WWP2-FL and -N isoform in the WWP2^−/−^ mouse kidneys (Supplementary Figure 3A).

We first examined the effect of WWP2 deficiency on UUO-induced renal fibrosis. After 14 days of UUO, Sirius red and Masson’s trichrome staining on the kidney sections revealed extensive renal fibrosis in WT mice (Figure 2A). This was significantly mitigated in WWP2^-/-^ mice, and was associated with a ∼50% reduction in the percentage of cortical fibrosis and decreased collagen deposition compared with UUO WT mice (Figure 2B). In line with the histological analysis, deletion of WWP2 attenuated the upregulation of ECM marker genes in UUO-induced kidneys at both protein and mRNA levels (Figure 2C-D). However, the effect of WWP2 deficiency on interstitial fibrosis was not apparent at 7-day UUO, a timepoint that coincides with mild-moderate renal fibrosis in WT animals (Supplementary Figure 3B-E).

**Figure 2.**
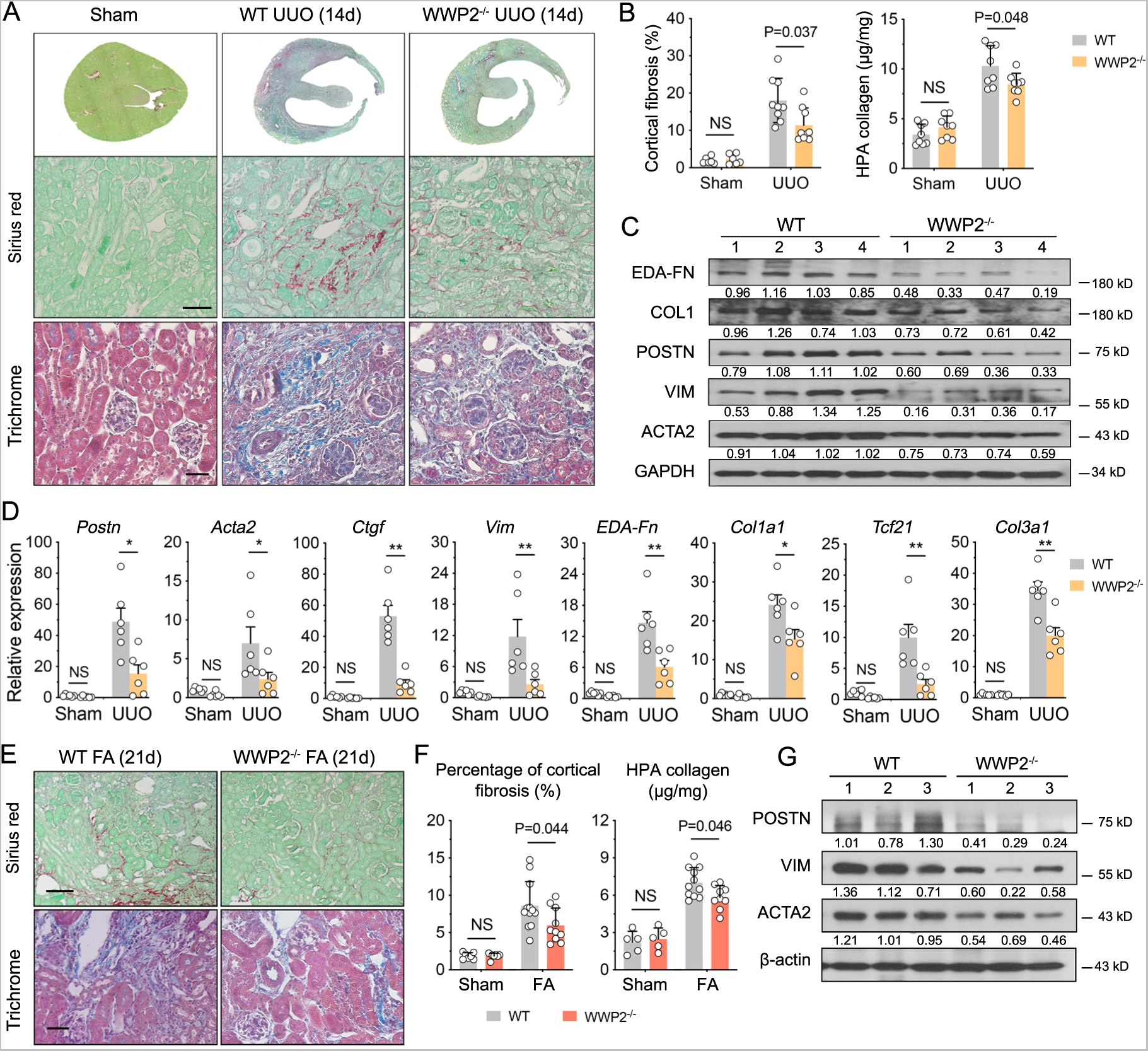
WWP2 deficiency protects from renal fibrosis *in vivo*. (A) Representative images of WT and WWP2^-/-^ mouse kidneys following UUO model for 14 days. (n=8 images recorded for each condition). *Top and middle panels*, Sirius Red staining for whole section and representative fibrotic area. Scale bars, 50 μm. *Bottom panels*, representative images of Masson’s trichrome staining for representative fibrotic area. Scale bars, 20 μm. (B) Quantitative analysis of cortical fibrosis-positive area (*left*, %) and HPA collagen levels (*right*, μg/mg) WT and WWP2^-/-^ mouse kidneys following UUO model for 14 days. n = 6-10, in each experimental group. (C) Representative western blot for ECM proteins in UUO kidney tissue from WT and WWP2^-/-^ mouse kidneys following UUO model for 14 days. (D) Expression of ECM genes, determined by RT-qPCR, in kidney from WT and WWP2^-/-^ mouse following UUO model for 14 days. (n=6, in each group). (E) Representative images of folic acid (FA)-induced fibrotic kidneys at day 21 in WT and WWP2^-/-^ mice (n=6 images recorded for each condition). *Top panel*, Sirius Red staining for representative fibrotic area. Scale bars, 50 μm. *Bottom panel*, representative images of Masson’s trichrome staining for representative fibrotic area. Scale bars, 20 μm. (F) Quantitative analysis of cortical fibrosis-positive area (*left*, %) and HPA collagen levels (*right*, μg/mg) in FA-induced fibrotic kidneys in WT and WWP2^-/-^ mice. n = 5-12, in each experimental group (G) Representative western blot for ECM proteins in FA-induced fibrotic kidneys from WT and WWP2^-/-^ mice. In each case, data values are reported as mean ± SD, and P-values were calculated by two-tailed Mann- Whitney U test.

We showed a protective effect of WWP2 deficiency in another widely used animal model of renal fibrosis, the folic acid-induced nephropathy (FA) model [45]. 21 days after FA induction, histochemistry for Sirius red and Masson’s trichrome showed reduced fibrotic lesions with lower percentage of cortical fibrosis and collagen content in WWP2^-/-^ when compared with WT controls (Figure 2E-F). The expression of ECM proteins in the kidney was also reduced in WWP2^−/−^ mice (Figure 2G).

These data from two *in vivo* fibrotic models demonstrate the protective effect of WWP2 deficiency towards renal fibrosis. Additional corroborative data in the transgenic WWP2 overexpressing mice (Supplementary Figure 2) further support a role of WWP2 in regulating renal fibrosis.

### WWP2 uncouples myofibroblast activation and proliferation in the fibrotic kidney

We previously demonstrated that WWP2 regulates heart fibrosis and cardiac fibroblast activation [40]. Here, we aimed to investigate the stromal cellular heterogeneity and the regulation of renal fibroblasts by WWP2 in kidney fibrosis. We carried out single-cell transcriptional analysis of fibrotic UUO kidneys, which identified two cell populations (clusters 7-8) functionally enriched for ECM organization and extracellular structure organisation (Supplementary Figure 4A-B). Using Pseudotime analysis we inferred the trajectory of gene expression changes in cells derived from UUO fibrotic kidney (Figure 3A, *left*), and confirmed that clusters 7-8 were characterized by a high ECM-expression score [46], including genes encoding for collagens, glycoproteins and proteoglycans (Figure 3A, *right*). In addition, clusters 7 and 8 were enriched for markers of myofibroblasts, and around 40% of cells expressed Acta2 (Supplementary Figure 4C). We thus defined these clusters 7-8 as renal myofibroblasts. WWP2 regulated the proportion of myofibroblasts in UUO-induced fibrotic kidneys. Specifically, we found a lower percentage of clusters 7-8 in WWP2^-/-^ mice compared with WT mice (Figure 3B), which was supported by flow cytometry analysis that showed a significantly reduced proportion of ACTA2^+^ cells following UUO (Figure 3C).

**Figure 3.**
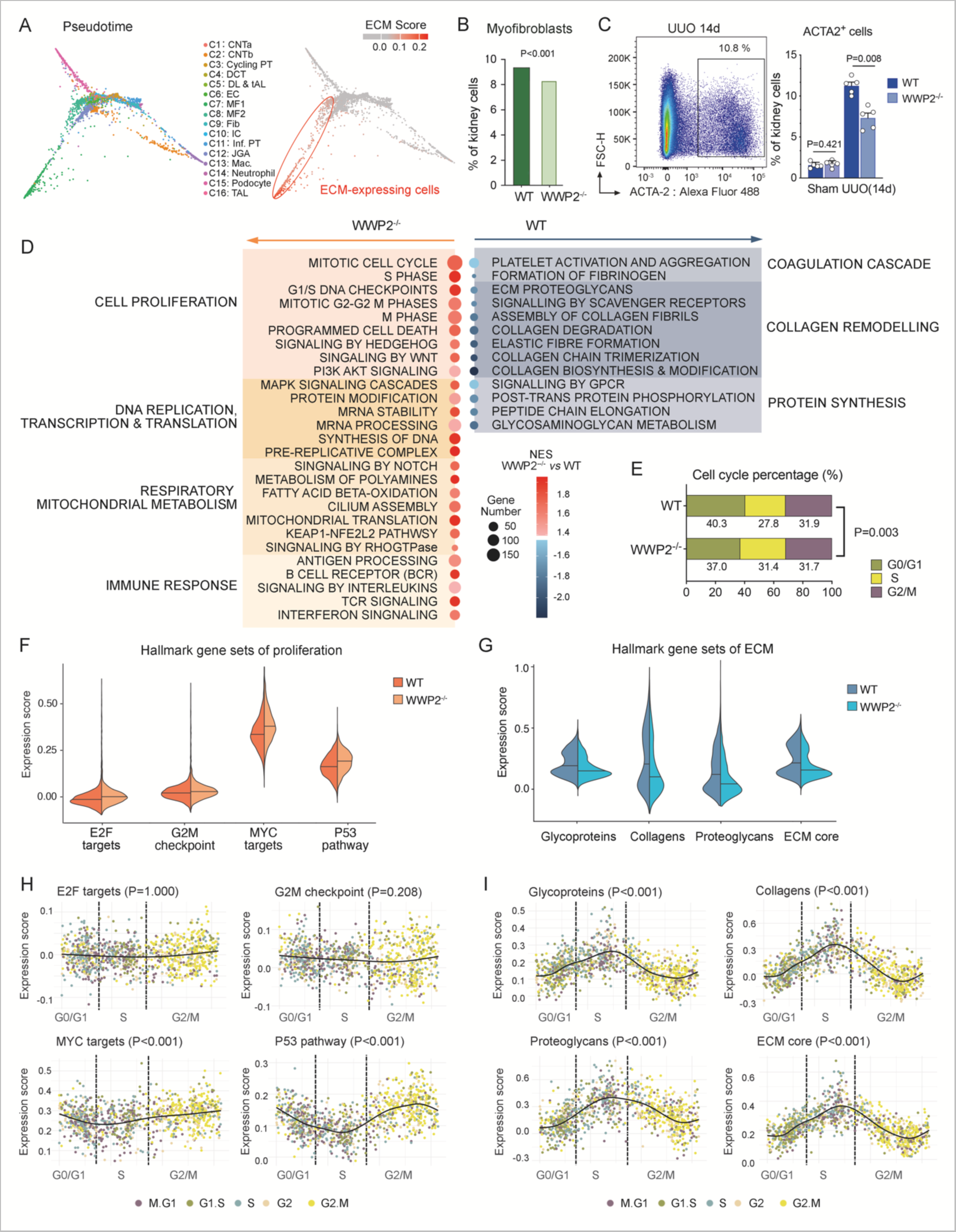
WWP2 mediates myofibroblasts phenotypes in fibrotic kidneys. (A) Diffusion map embedding of single cells data from UUO kidneys (*left*), and relative expression of ECM genes over imposed on the same embedding (*right*). Compared with other cell types, two myofibroblasts clusters (C7 and C8) presented high expression scores of ECM genes, quantified by a composite “ECM score” (see Methods). (B) Bar plots showing the percentage of myofibroblasts derived from UUO kidneys in WT and WWP2^-/-^ mice. Myofibroblasts percentage calculated with respect to all renal cells, and P-value was calculated by chi-square (χ^2^) test (df=1) for cell proportions. (C) *Left*, representative graph of ACTA2^+^ cells in WT mouse using flow cytometry in renal living cells from UUO kidney (14 days). *Right*, quantification of ACTA2+ cells in kidney cells from WT and WWP2^-/-^ mice (n=5, from 3 independent experiments). Values are reported as mean ± SD and P-values calculated by two-tailed Mann-Whitney U test. (D) Classification of Reactome pathways enriched in ECM-expressing myofibroblasts and significantly different between WT and WWP2^-/-^ UUO kidneys by Gene Set Enrichment Analysis (GSEA) (false discovery rate (FDR) <0.05). NES, normalized enrichment score, where a positive NES indicate upregulation in WWP2^-/-^ compared with WT myofibroblasts. (E) Proportions of ECM-expressing myofibroblasts at different phases of the cell cycle, grouped as G0/G1, S and G2/M phases. ECM-expressing myofibroblasts from UUO kidneys are grouped according to WT and WWP2^-/-^ genotypes. P-value was calculated by chi-square (χ^2^) test for cell number in G0/G1, S and G2/M phases, yielding χ^2^ = 11.89, df=2, p-value = 0.003 (F-G) Expression score for hallmark gene sets for proliferation (panel F) and ECM production genes (panel G) in ECM-expressing myofibroblasts from UUO fibrotic kidneys. See Methods for definition of gene sets and score calculation. For each gene set, difference in expression score between WT and WWP2^-/-^ groups was tested using the non-parametric Wilcoxon rank-sum test; for each given gene set the P-value for the difference was <0.001 (H-I) Using the Revelio algorithm (see Methods), myofibroblast cells derived from 5 human CKD kidneys [11] are arranged along a pseudotime trajectory based on to their cell cycle phases and grouped as G0/G1, S, and G2/M. The distribution of hallmark gene set scores for proliferation (panel H) and ECM production (panel I) is shown for each myofibroblasts arranged accordingly to its cell cycle phase and the main trend is approximated by smooth line interpolation (bold black line). For each gene set, P-values for significance of change in the linear trend (bold black line) were calculated by local regression-based WAVK test (see Methods for additional details).

We used Gene-Set Enrichment Analysis (GSEA) to map the cellular processes and pathways modulated by WWP2 (Figure 3D). Renal myofibroblasts from WWP2^-/-^ UUO mice showed increased expression of genes related to the mitotic cell cycle, cell division phases, and DNA replication, transcription, and translation, which were previously reported to contribute to cellular proliferation [47]. In keeping with this, WWP2^-/-^ myofibroblasts had a lower proportion of cells in G0/G1 phases of the cell cycle compared with WT mice in fibrotic kidneys (Figure 3E), and a higher expression of the hallmark gene sets for cell proliferation, including MYC targets and P53 pathways (P<0.001 for each gene set, Figure 3F). In contrast, the expression of canonical pathways associated with ECM production, particularly collagen remodeling, was significantly reduced in WWP2^-/-^ myofibroblasts (Figure 3D). The hallmark gene sets for core ECM matrisome were also downregulated in WWP2^-/-^ myofibroblasts compared to WT cells (P<0.001 for each gene set, Figure 3G). Therefore, WWP2 differentially regulates cellular proliferation and pro-fibrotic cell activation, suggesting these processes are inversely correlated in renal myofibroblasts.

Similarly, in myofibroblasts from human CKD kidneys we observed that cellular proliferation and pro-fibrotic activation were differentially active across cell cycle phases. Using the Revelio algorithm (see Methods), we extracted cell cycle information for renal myofibroblasts derived from 10 CKD patients [11], and investigated the dynamic expression changes in cell proliferation and ECM processes. The MYC targets and P53 pathway gene sets exhibited dynamic transcriptional changes, with the highest expression observed at G2/M phase (P<0.001, Figure 3H), while all ECM gene sets showed the lowest expression at G2/M phase (Figure 3I). Thus, single cell analysis in CKD kidneys suggests myofibroblast heterogeneity, with the proliferation and pro-fibrotic activation processes exhibiting different dynamics across cell cycle phases.

These analyses in mice and human renal myofibroblasts indicate that cell proliferation and pro-fibrotic activation might represent different states of the same myofibroblast subpopulation, and these states respond transcriptionally to different cell cycle stages. Both these states are regulated by WWP2, which attenuates fibrosis by reducing myofibroblasts formation and also regulating their function, promoting cellular proliferation while inhibiting collagen synthesis, respectively. Furthermore, upregulation of pathways associated with respiratory mitochondrial metabolism in WWP2^-/-^ myofibroblasts (Figure 3D) suggested a potential metabolic regulatory role of WWP2 in fibrotic myofibroblasts.

### WWP2 promotes fibrotic activation and suppresses proliferation of myofibroblasts

The regulation of myofibroblasts by WWP2 was further characterized *in vitro* by culturing primary renal fibroblasts isolated from WT and WWP2^-/-^ mice. The WWP2^-/-^ cells lack WWP2 expression (Figure 4A), and TGFβ1 treatment upregulates WWP2 during myofibroblasts differentiation (Supplementary Figure 5A). Flow cytometry analysis showed that the majority of cultured WT myofibroblasts expressed ACTA2, and the fraction of ACTA2-expressing cells was reduced in WWP2^-/-^ myofibroblasts (Figure 4B).

**Figure 4.**
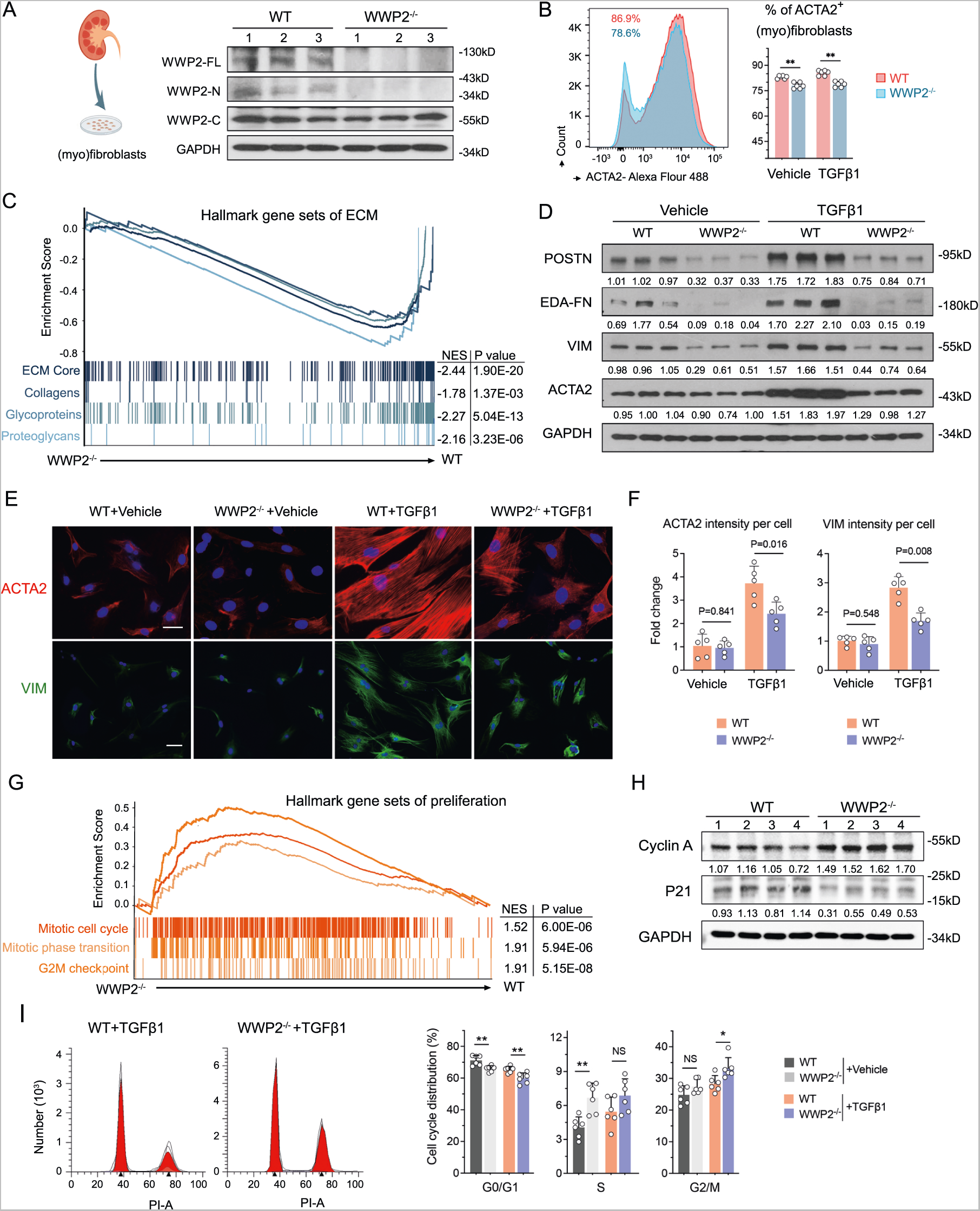
WWP2 deficiency promotes cellular proliferation and supresses pro-fibrotic activation in renal myofibroblasts in vitro. (A) Representative western blot showing the levels of WWP2 in primary cultured renal myofibroblasts (P2) derived from WT and WWP2^-/-^ kidneys. WWP2^-/-^ cells lack WWP2-full length and -N isoforms expression. (B) *Left*: representative graph of ACTA2^+^ cells in cultured renal myofibroblasts (P2) derived from WT and WWP2^-/-^ mice using flow cytometry. *Right*: quantification of ACTA2+ cells in cultured myofibroblasts derived from WT and WWP2^-/-^ kidneys (n = 6, from 3 independent experiments). Values are reported as mean ± SD. TGFβ1 (5ng/μl) for 72 hours. (C) Gene Set Enrichment Analysis (GSEA) of ECM pathways in cultured renal myofibroblasts, treated with TGFβ1 (5ng/μl) for 72 hours, showing the Enrichment Score for ECM gene sets in WWP2^-/-^ (x-axis, left) and WT kidneys (x-axis, right). NES, normalized enrichment score, where negative values indicate downregulation of the gene set in WWP2^-/-^ myofibroblasts with respect to WT myofibroblasts. (D) Representative western blotting for ECM proteins in cultured TGFβ1-treated myofibroblasts (P2) derived from WT and WWP2^-/-^ kidneys. TGFβ1 (5ng/μl) for 72 hours. (E-F) Representative microscopy images (E) and quantification analysis (F) with immunostaining for ACTA2 and VIM in cultured renal TGFβ1-treated myofibroblasts derived from WT and WWP2^-/-^ kidneys (n=5 biological replicates). One single dot indicates the average of 25-40 myofibroblasts taken from each slide. Values are reported as mean ± SD. TGFβ1 (5ng/μl) for 72 hours (G) Similar to the data in panel (C), GSEA of proliferation pathways in cultured renal myofibroblasts (P2) treated with TGFβ1 (5ng/μl) for 72 hours. Enrichment Score comparing WWP2^-/-^ and WT myofibroblasts. Positive NES values indicate upregulation in WWP2^-/-^ myofibroblasts with respect to WT myofibroblasts. (H) Representative western blotting for Cyclin A and P21 in cultured TGFβ1-treated renal myofibroblasts (P2) derived from WT and WWP2^-/-^ kidneys. TGFβ1 (5ng/μl) for 24 hours (I) *Left*: representative graph of cell cycle in cultured TGFβ1-treated renal myofibroblasts (P2) derived from WT and WWP2^-/-^ mice using flow cytometry. *Right*: quantification of cell cycle at G0/G1, S and G2/M phases in cultured myofibroblasts derived from WT and WWP2^-/-^ kidneys (n = 6, from 3 independent experiments). TGFβ1 (5ng/μl) for 24 hours. Values are reported as mean ± SD.

We performed bulk RNA sequencing and molecular analysis to investigate the role of WWP2 in myofibroblast response to TGFβ1 treatment *in vitro*. In keeping with the scRNA-seq analysis of the myofibroblasts derived from UUO kidneys (Figure 3G), GSEA showed that cultured WWP2^-/-^ myofibroblasts exhibit a significantly decreased expression of gene sets related to the ECM compared with WT myofibroblasts (Figure 4C). The increased expression of fibrous proteins, such as ACTA2, EDA- FN and POSTN, which was observed upon TGFβ1 treatment, was largely diminished in WWP2^-/-^ myofibroblasts (Figure 4D). Furthermore, TGFβ1-stimulated WT myofibroblasts presented a clear organization of ACTA2 into stress fibers, while WWP2^-/-^ myofibroblasts showed a diffuse expression of ACTA2 and VIM with rare incorporation into stress fibers (Figure 4E). We confirmed a significantly lower ACTA2 and VIM intensity per cell in WWP2^-/-^ myofibroblasts compared to WT cells (Figure 4F).

Confirming our previous results (Figure 3F), GSEA showed that gene sets related to cell proliferation were more prominently expressed in WWP2^-/-^ myofibroblasts compared with WT (Figure 4G). The expression level of cell division phase checkpoint proteins P21 and Cyclin A in cultured myofibroblasts was also regulated by WWP2 (Figure 4H). Cell cycle analysis showed that the majority of cultured renal myofibroblasts were in G0/G1 phase (∼70%), which was slightly decreased after TGFβ1 treatment and further decreased upon deletion of WWP2^-/-^, while cells in S or G2/M phase showed the opposite trend (Figure 4I).

Corroborating our previous observations, we showed that WWP2 promoted fibrotic activation and suppressed proliferation of myofibroblasts in cells with WWP2 overexpression (WWP2^OE^) (Supplementary Figure 5B). WWP2^OE^ led to a further enhanced production of Fibronectin 1 in myofibroblasts upon TGFβ1 treatment (Supplementary Figure 5C), and a lower percentage of myofibroblasts in G2/M phase (Supplementary Figure 5D).

### WWP2 regulates myofibroblast energy metabolism during fibrosis

Our single cell analysis in UUO kidneys suggested a potential metabolic regulatory role of WWP2 in fibrotic myofibroblasts (Figure 3D). To investigate this in detail, we first used Compass [48] to profile the metabolic reactions in renal myofibroblasts from UUO-mouse and human CKD patients. In fibrotic myofibroblasts derived from mice, WWP2 deficiency yielded increased activity in a range of metabolic reactions (Supplementary Figure 6A), especially those related to central carbon metabolism and amino acid metabolism (Figure 5A). These data suggest that WWP2 deficiency is associated with higher glucose utilization, including glucose utilisation in the pentose phosphate pathway (PPP), and TCA cycle. WWP2 deficiency increases fatty acid oxidation (FAO), another pathway that induces TCA cycle activity and mitochondrial respiratory metabolism [49, 50]. WWP2 also influences amino acid metabolism, which contributes to the balance of the NAD(P)^+^/NAD(P)H redox pair, serving as an extra metabolic energy source [51].

**Figure 5.**
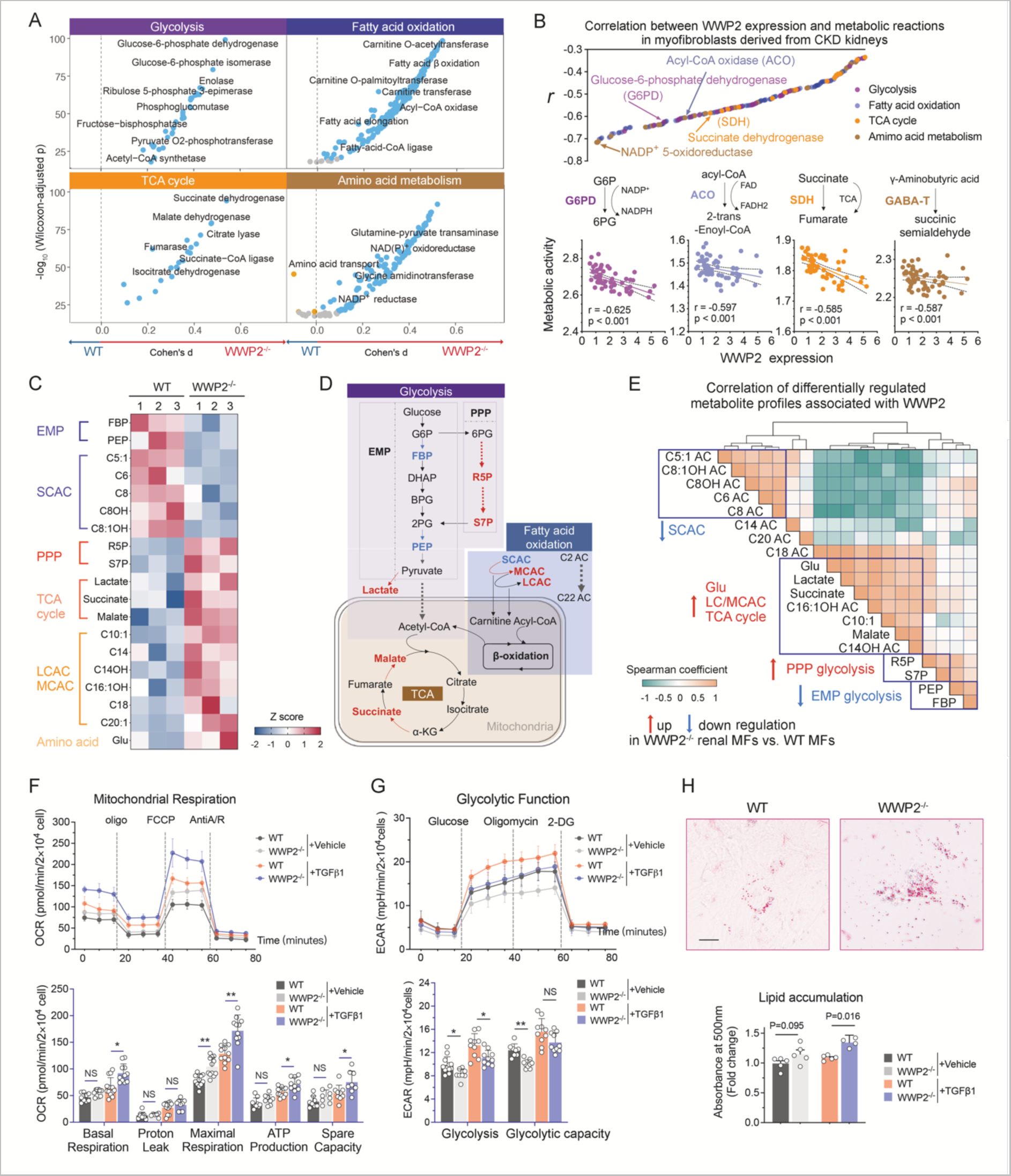
Differential energy metabolism of WT and WWP2^-/-^ myofibroblasts. (A) Compass-score differential activity test in ECM-expressing myofibroblasts derived from WT and WWP2^-/-^ UUO kidneys, for reactions in the glycolysis, TCA cycle, fatty acid oxidation and amino acid metabolism pathways. Statistical significance (y-axis) for the difference in the activities scores of metabolic resections between WWP2^-/-^ and WT group was assessed by non-parametric Wilcoxon rank-sum test, while the effect size was estimated by Cohen’s D (x-axis). The whole set of metabolic reactions changes in myofibroblasts are in Supplementary Figure 6A. (B) WWP2 mRNA expression is negatively associated with activity of metabolic resections in renal myofibroblasts from 5 human CKD patients’ kidneys [11]. *Upper panel*: significant Spearman correlations (P<0.05) between Compass scores of metabolic resections and the expression level of WWP2 in renal myofibroblasts. The color coding represents different metabolic pathways of the metabolic reactions. *Lower panels*: detail on the correlation between WWP2 expression and activity of 4 key metabolic reactions. The data for whole set of metabolic reactions is showed in supplementary Figure 6B. (C) Metabolomics analysis shows the profile of metabolites exhibiting significant differences (P<0.05, false discovery rate (FDR)<17%, two-tailed non-parametric Mann-Whitney U test) between cultured TGFβ1-treated (5ng/μl for 72 hours) renal myofibroblasts (P2) derived from WT and WWP2^-/-^ kidneys. Each metabolite level was normalized and represented as Z score. EMP, Embden-Meyerhof pathway of glycolysis; PPP, pentose phosphate pathway; SCAC, small chain of acylcarnitine; LCAC, long chain of acylcarnitine; MCAC, medium chain of acylcarnitine; FBP, fructose-1,6-bisphosphate; PEP, phosphoenolpyruvate; R5P, ribose-5-phosphate; S7P: Sedoheptulose-7-phosphate; Glu: glutamic acid. (D) Simplified overview of central metabolic fluxes in glycolysis, TCA cycle, fatty acid oxidation and amino acid metabolism in cultured renal myofibroblasts derived from WT and WWP2^-/-^ kidneys. Colour indicates the relative metabolite or enzyme rates in WWP2^-/-^ myofibroblasts compared to WT cells, where red indicates higher flux in WWP2^-/-^ myofibroblasts and blue indicates higher flux in WT cells. G6P, Glucose-6-phosphate; DHAP, Dihydroxyacetone phosphate; BPG, Bisphosphoglycerate or 2,3-Bisphosphoglycerate; 2PG, 2-Phosphoglycerate; α-KG, a-ketoglutarate. (E) Coordinated regulation of metabolites associated with WWP2 deficiency. Metabolite-metabolite correlation analysis of differential metabolites (reported in panel C). Positive correlations are depicted in orange, while negative correlations are represented in green. Metabolites that exhibit a high degree of correlation are highlighted (blue rectangles). (F) *Upper panel:* representative Seahorse Mito stress assays for the Oxygen Consumption Rate (OCR) in cultured TGFβ1-treated (5ng/μl for 72 hours) renal myofibroblasts derived from WT and WWP2^-/-^ kidneys. *Lower panel:* barplots summarizing the phenotypes derived from OCR analysis results obtained from three independent biological experiments. Each experiment involved independent cell plating on 3-4 Seahorse microplate wells. (G) *Upper panel*: representative Seahorse Glycolysis assays for the Extra Cellular Acidification Rate (ECAR) in cultured renal myofibroblasts derived from WT and WWP2^-/-^ kidneys. *Lower panel*: barplots summarizing the phenotypes derived from ECAR analysis from three independent biological experiments. Each experiment involved independent cell plating on 3-4 Seahorse microplate wells. (H) Visualization (*upper panels*) and quantification (*lower panels*) of neutral lipids by oil red O analysis in cultured TGFβ1-treated (5ng/μl for 72 hours) renal myofibroblasts derived from WT and WWP2^-/-^ kidneys. Scale bars, 50 μm. Quantification was obtained from four independent experiments and data values are reported as mean ± SD. P-values were calculated by two-tailed Mann-Whitney U test.

The changes in metabolic reactions mediated by WWP2 was corroborated by the analysis of renal myofibroblasts derived from CKD patients (see Methods and Supplementary Figures 6B). In these myofibroblasts, WWP2 expression was negatively correlated with reactions necessary for mitochondrial energy metabolism (Figure 5B; *top panel*). Specifically, our findings underscore several reactions (Figure 5B; *bottom panels*), including glucose-6-phosphate dehydrogenase (G6PD) catalyzed reaction, the first step in the PPP [52] which leads to the conversion of NADP to NADPH [53]. We also highlight Acyl-CoA oxidase (ACO), responsible for the oxidation of fatty acids, acting on CoA derivatives of fatty acids with aliphatic carbons between medium to long chain [54], as well as 4-aminobutyrate transaminase (GABA-T) participating glutamate metabolism and tightly coupled to cellular energy metabolism [55], and succinate dehydrogenase (SDH) as part of the TCA cycle [56]. The association of WWP2 and these enzymatic reactions linked with mitochondrial respiration, was confirmed in a second CKD cohort (see Methods and Supplementary Figure 7A-B). Notably, we found a stronger negative association between WWP2 expression and these metabolic reactions in CKD patients compared to controls (Supplementary Figure 7C), suggesting that the downregulation of these metabolic reactions by WWP2 may be more pronounced in disease.

We then took a metabolomics approach to detail the response of the cellular energy metabolism in WT and WWP2^-/-^ renal myofibroblasts treated with TGFβ1 (Supplementary Figure 8 and Figure 5C-D). WWP2^-/-^ myofibroblasts showed increased ribose-5-phosphate (R5P) and sedoheptulose-7-phosphate (S7P) in the PPP, at the expense of fructose 1,6-bisphosphate (FBP) and phosphoenolpyruvate (PEP) in the Embden-Meyerhof-Parnas (EMP) pathway. WWP2^-/-^ myofibroblasts also showed changes in acylcarnitine (AC) facilitating FAO for energy production in mitochondria [57], including an increase in long/medium chain AC (LC/MCAC) and a decrease in small chain AC (SCAC). The level of glutamate (Glu) was higher in WWP2^-/-^ myofibroblasts compared with WT cells. In addition, we found increased levels of lactate, malate, and succinate in WWP2^-/-^ myofibroblasts, indicative of an enhanced TCA cycle [56, 58], thus confirming the results of COMPASS (which were based on single cell data analysis). These metabolites were co-regulated by WWP2 (Figure 5E), which might imply a synergistic impact of WWP2 deletion on multiple interconnected metabolic pathways.

Given that TCA cycle is coupled with OXPHOS to produce ATP, we next sought to evaluate mitochondrial function by Seahorse analysis (Figure 5F-G). Compared with WT cells, WWP2^-/-^ myofibroblasts showed a higher oxygen consumption rate (OCR), with increased levels of basic respiration, maximum respiration and ATP production (Figure 5F), and exhibited a decreased extracellular acidification rate (ECAR) (Figure 5G). These results confirmed the increased FAO and TCA cycle activity observed following metabolomics analysis (Figure 5C). Interestingly, WWP2^-/-^ myofibroblasts also show significantly higher lipid droplet accumulation, as indicated by Oil Red O staining (Figure 5H), suggesting that FAO may be associated with an increased uptake of fatty acids stored in lipid droplets

Taken together, these data demonstrate a role for WWP2 in regulating components of myofibroblast energy metabolism (Supplementary Figure 8E). These data would be consistent with WWP2 deficiency boosting glucose utilization through the PPP concomitant with FAO leading to enhanced TCA cycle and ATP production. This metabolic rewiring in myofibroblasts might underlie changes in cell proliferation and ECM production observed in WWP2^-/-^ cells (Figure 4) and in kidney fibrosis (Figures 2-3).

### PGC-1α signalling mediates the WWP2-regulated myofibroblasts metabolism

WWP2 has been reported to act as a co-factor in nuclear transcriptional complexes [40, 59], and we show that its localization changes upon TGFβ1 treatment within the nucleus (Supplementary Figure 9a). We previously demonstrated that WWP2 regulates transcription factor activity of SMAD2 in cardiac fibroblast [40] and of IRF7 in cardiac macrophages [41]. Therefore, we hypothesise that WWP2 plays a similar transcriptional regulatory role in fibrotic renal myofibroblasts. Given the conserved regulatory role of WWP2 in FAO during murine and human renal fibrosis, we focused on PGC-1α, a critical modulator of FAO [28, 29], which has been previously shown to regulate myofibroblast metabolism during tissue fibrosis [32–34]. To investigate the potential transcriptional regulation of PGC-1α by WWP2, we first used ChIP-Seq analysis and assessed WWP2 binding on the *Ppargc1a* locus, which encodes for PGC-1α (Figure 6A). We further designed primers (grey boxes, Figure 6A) for ChIP-qPCR near the transcription start site (TSS) of the *Ppargc1a* locus. Specially, genomics regions I-III contain WWP2 binding signals (ChIP-Seq peaks, in red) and known DNA-binding elements (light yellow boxes) identified from the REMAP DB [60], while regions IV to VI served as controls. Upon TGFβ1 stimulation, there was a significant increase in the binding of the WWP2 transcriptional complex to the *Ppargc1a* locus (regions I to III) compared to the control groups (regions IV to VI) (Figure 6B). WWP2 deficiency was associated with increased expression of PGC-1α in cultured myofibroblasts at protein and mRNA levels (Figure 6C and Supplementary Figure 9B). This analysis suggests that WWP2 may inhibit the transcription of PGC- 1α in activated myofibroblasts.

**Figure 6.**
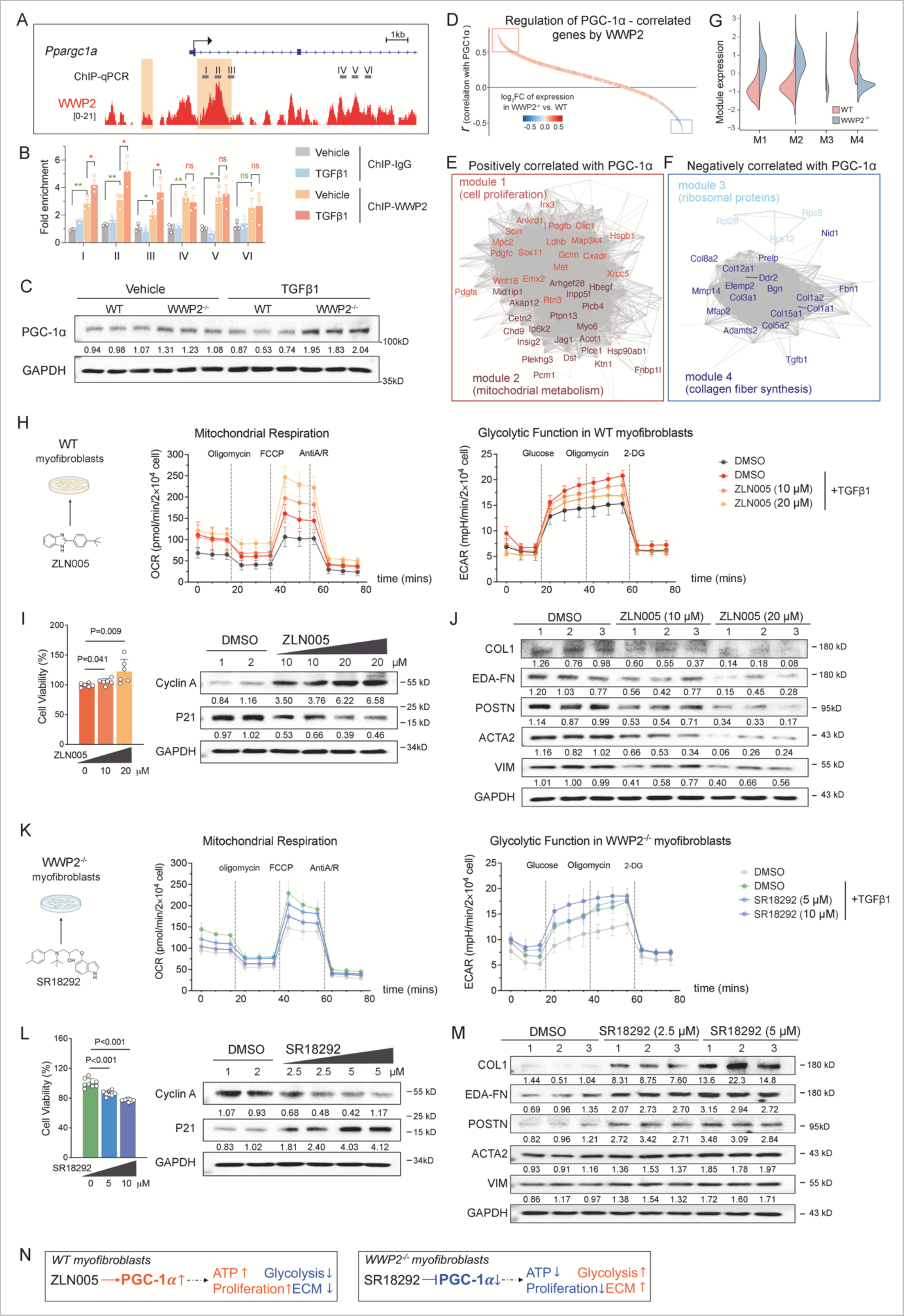
WWP2 regulates renal myofibroblasts via PGC-1α-mediated mitochondrial respiratory function. (A) Distribution of WWP2 ChIP-seq signal (red) at the *Ppargc1a* locus in primary WT myofibroblasts after TGFβ1 stimulation (5ng/μl, 72 hrs). The primers designed for ChIP-qPCR (grey boxes, I-VI) and the reference *Ppargc1a* gene (blue) are indicated. The yellow boxes represent genomic sequence locations with *cis*-regulatory elements around the WWP2 peaks identified by ChIP-seq analysis, predicted by Regulatory Sequence Analysis Tools (RSAT) (see Methods for details). (B) ChIP-qPCR analysis shows increased binding of WWP2 at *Ppargc1a* genomic regions I, II and III after TGFβ1 stimulation, which overlap with predicted *cis*-regulatory elements. Each bar shows WWP2 enrichment normalized to input and the ChIP-IgG control. Values are reported as mean ± SD (n = 3, from 3 independent experiments). P-values were calculated by two-tailed Mann-Whitney U test. *, P<0.05; **, P<0.01; ns, P>0.05. (C) Representative western blot for PGC-1α in cultured renal myofibroblasts derived from WT and WWP2^-/-^ kidneys. TGFβ1 stimulation (5ng/μl) for 24 hrs. (D) Genome-wide distribution of genes sorted based on their Pearson correlation with PGC-1α expression in myofibroblasts derived from fibrotic mouse kidneys. The colour scale represents the fold change (log_2_FC) in expression between WWP2^-/-^ myofibroblasts compared to WT cells. The red rectangle includes genes positively related to PGC-1α (*r* > 0.5, P<0.001) and which are upregulated in WWP2s^-/-^ myofibroblasts compared to WT cells; the genes in the blue rectangle are negatively related to PGC-1α (*r* < −0.5, P<0.001) and downregulated in WWP2^-/-^ myofibroblasts. (E-F) Subset of network genes from those highlighted in the red (D) and blue (E) rectangles in panel C, which are correlated with PGC-1α (|*r|* > 0.6). PGC-1α-network genes are further clustered in 4 distinct modules (M1-M4), positively (M1-M2) and negatively (M3-M4) correlated with PGC-1α, respectively (see Methods for details). Within each network, edges represent positive co-expression relationships between genes, and colours indicate functional clustering of the module genes, highlighting the cell proliferation (M1) and mitochondrial metabolism (M2) (panel D), and processes related to fibrous protein synthesis (M3 and M4, panel E). The genes highlighted are more strongly correlated positively (r > 0.8, panel E) or negatively (r < −0.8, panel F) with PGC-1α expression, respectively, in myofibroblasts for fibrotic mouse kidneys. (G) For each module (M1-M4, see panels E-F), violin plot shows the distribution of normalized gene expression levels derived from bulk-RNA seq analysis of cultured WT and WWP2^-/-^ myofibroblasts treated with TGFβ1 (5ng/μl, 72 hrs). For each module, P-value for the difference between WT and WWP2^-/-^ myofibroblasts was <0.001, which was calculated by two-tailed Wilcoxon rank-sum test. (H) Effects of ZLN005 (10 μM and 20 μM) on cell metabolism in cultured WT myofibroblasts after TGFβ1 treatment (5ng/μl, 72 hrs), measured by Searhorse assay. DMSO (10 μM) was used as a reference control. *Left panel*: experimental design schematic. *Middle panel*: representative Seahorse Mito stress assays for OCR. *Right panel*: representative Seahorse Glycolysis assays for ECAR. Results of quantification analysis are shown in Supplementary Figure 9D. (I) Effects of ZLN005 (10 μM and 20 μM) on cell proliferation in cultured WT myofibroblasts after TGFβ1 treatment (5ng/μl, 72 hrs), measured by MTT assay (*left panel*) and western blot (*right panel*). P-values were calculated by two-tailed Mann-Whitney U test (J) Western blot showing the effect of ZLN005 (10 μM and 20 μM) on ECM proteins production in cultured WT myofibroblasts after TGFβ1 treatment (5ng/μl, 72 hrs). (K) Effects of SR18292 (2.5 μM and 5 μM) on cell metabolism in cultured WWP2^-/-^ myofibroblasts after TGFβ1 treatment (5ng/μl, 72 hrs) by Searhorse assay. DMSO (5 μM) was used as a reference control. *Left panel*: experimental design schematic. *Middle panel:* representative Seahorse Mito stress assays for OCR. *Right panel:* representative Seahorse Glycolysis assays for ECAR. Results of quantification analysis are shown in Supplementary Figure 9E. (L) Effects of SR18292 (2.5 μM and 5 μM) on cell proliferation in cultured WWP2^-/-^ myofibroblasts after TGFβ1 treatment (5ng/μl, 72 hrs), measured by MTT assay (*left panel*) and western blot (*right panel*). P-values were calculated by two-tailed Mann-Whitney U test (M) Western blot showing the effect of SR18292 (2.5 μM and 5 μM) on ECM proteins production in cultured WWP2^-/-^ myofibroblasts after TGFβ1 treatment (5ng/μl, 72 hrs), measured by Western blot. (N) Schematic summary of the cellular and metabolic phenotypes observed following PGC-1α pharmacological activation (by ZLN005) or inhibition (by SR18292) in primary cultured WT and WWP2^-/-^ renal myofibroblasts, respectively.

To further investigate the role of WWP2 on PGC-1α in myofibroblasts we interrogated the possible transcriptional activation of the PGC-1α regulatory network using single cell data from the fibrotic mouse kidney. We first derived the sets of genes co-expressed with *Ppargc1a* in myofibroblasts (see Methods), and then queried the transcriptional effect of WWP2 deletion (Figure 6D). The genes positively correlated with *Ppargc1a* were found to be upregulated in WWP2^-/-^ myofibroblasts (red box, Figure 6D); and were enriched for cell proliferation (module 1) and mitochondrial metabolism (module 3, Figure 6E). In contrast, genes negatively correlated with *Ppargc1a* were downregulated in WWP2^-/-^ myofibroblasts (blue box, Figure 6D), and these genes contribute to collagen synthesis, including ribosomal protein genes (module 3) and collagen fiber genes (module 4, Figure 6F). Bulk RNA-seq analysis of TGFβ1-treated myofibroblasts confirmed the upregulation of modules 1-2 and the downregulation of modules 3-4 in response to WWP2 deficiency (Figure 6G). These results suggest a role for WWP2 in regulating PGC-1α and its regulatory transcriptional network in myofibroblasts, which in turn is important for mitochondrial function, cell proliferation and pro-fibrotic activation. We postulate that the WWP2-PGC- 1α-axis might contribute to the observed metabolic phenotypes regulated by WWP2 in myofibroblasts (Supplementary Figure 8E).

However, it remains unclear whether PGC-1α mediates the metabolic phenotypes regulated by WWP2 in myofibroblasts. To explore this, we employed an activator (ZLN005) and inhibitor (SR18292) of PGC- 1α in primary cultured WT and WWP2^-/-^ myofibroblasts. In WT myofibroblasts treated by TGFβ1, ZLN005 upregulated PGC-1α expression in a dose-dependent manner (Supplementary Figure 9C), leading to increased mitochondrial respiration and decreased glycolysis, as evidenced by elevated OCR levels and reduced ECAR levels, respectively (Figure 6H and Supplementary Figure 9D). Furthermore, ZLN005 treatment enhanced myofibroblast cell proliferation (Figure 6I) and consistently, reduced pro-fibrotic markers in myofibroblast (Figure 6J). In contrast, SR18292 treatment reversed the metabolic response in WWP2-deficient myofibroblasts (Figure 6K and Supplementary Figure 9E-F), which was mirrored by cell proliferation arrest (Figure 6L) and pro-fibrotic proteins production (Figure 6M). In summary, these experiments show that WWP2 modulates upstream the transcriptional activity of PGC-1α, a master regulator in mitochondria energy metabolism, particularly FAO [61] and PPP [62]. They also suggest that myofibroblast proliferation and ECM production are linked to the metabolic status of the renal myofibroblast.

## Discussion

Fibrogenesis is major driver of CKD progression, which is meditated by several renal cell types. Here, we took a myofibroblast-centric view on kidney tubulointerstitial fibrosis, and we highlight the biological relevance of cell metabolism for myofibroblasts proliferation and pro-fibrotic activation in CKD progression. Fibrosis is not limited to kidney diseases, but is also associated with chronic progressive failure in various organs such as the liver, lung, heart, intestine, and skin [63]. We show that myofibroblasts function in renal fibrosis is dependent on a WWP2-mediated metabolic regulation. WWP2 has previously been implicated in cardiac fibrosis where it acts as a regulator of ECM network genes [40]; our data in human and experimental models of CKD suggest a possible wider role of WWP2 in fibrosis in organs other than the heart.

In our study, we report the association between the expression level of WWP2 and the severity of tubulointerstitial fibrosis, as well as kidney dysfunction in CKD patients and mouse models of kidney fibrosis. Notably, deficiency of WWP2 yielded a protective effect against fibrotic progression in the kidneys, whereas overexpression of WWP2 exacerbated kidney fibrosis. *In vitro* overexpression of WWP2 has been recently proposed to ameliorate apoptosis of IR-induced HK-2 cells, an acute kidney injury model [64]. However, the latter was a study based on cell lines, which might not reflect the function of WWP2 in renal pathophysiology.

Myofibroblasts are the main source of ECM and play a significant role in the development of tissue fibrosis [4]. We used scRNA-seq to account for cellular heterogeneity and identify multiple phenotypes for renal myofibroblasts in fibrotic mouse kidney. By doing so, we aim to reveal functionally specialized myofibroblast populations and their potential regulation by WWP2. During kidney fibrosis, myofibroblasts exhibit a dual phenotype: cell proliferation and pro-fibrotic activation [65], both significant contributors to the net ECM production in the tubulointerstitium and subsequent progression of CKD [18, 25, 66, 67]. The activated pro-fibrotic myofibroblasts are responsible for synthesizing fibrous proteins, which are the primary source of ECM production in fibrotic kidneys [68]. Myofibroblasts have the capacity to proliferate in response to cytokine cues, which lead to expansion of the fibroblast population in the diseased renal interstitial space [13, 69]. Thus, our data support the model whereby pro-fibrotic activation and proliferation of myofibroblasts are linked and the fine balance between the two processes determine the rate of ECM production and the severity of renal fibrosis [13, 14]. Indeed, we show a decline in the expression of pro-fibrotic activation genes associated with the transition from G0/G1 to G2/M during the cell cycle dynamics of myofibroblasts. In contrast, the cell proliferation process showed an increase during the cell cycle dynamics of myofibroblasts. Lemons et al. [20] also reported that myofibroblasts arrested in their cell cycle, exhibit higher expression of fibrous proteins. Notably, WWP2 uncouples these two phenotypes of myofibroblasts acting as an upstream regulator: WWP2 deficiency suppresses the pro-fibrotic activation and enhances cell proliferation in both fibrotic kidneys and cultured myofibroblasts.

During fibrotic activation and cell proliferation, myofibroblasts exhibit high metabolic activity [20]. Intriguingly, WWP2 regulates metabolic pathways in myofibroblasts, particularly those involved in mitochondrial respiratory function. In two independent CKD patient cohorts, the expression of WWP2 expression negatively associated with the activity of key metabolic reactions in myofibroblasts. This negative association was confirmed by genetic deletion of WWP2 *in vivo*, where myofibroblasts derived from WWP2^-/-^ kidneys exhibited enhanced metabolic rates in central carbon metabolism and amino acid pathways. Within these pathways, we found important metabolic fluxes being increased in WWP2- deficient myofibroblasts. In particular, succinate, and LCAC actively participate in the processes of fatty acid oxidation and the TCA cycle, which directly contribute to mitochondrial function in cells [70]. The increased ATP production in WWP2-deficient myofibroblasts supports the activation of fatty acid oxidation which correlates with increased TCA cycle activity. In keeping with our findings, renal fibrosis is frequently characterized by a decrease in mitochondrial biogenesis [71] and damage to mitochondrial DNA (mtDNA) [72]. Furthermore, the protective WWP2 deficiency resulted in a reduction in cell glycolysis, particularly through the EMP pathway, while metabolic fluxes toward the PPP were increased. The regulation of this pathway by WWP2 is also important, as inhibition of glycolysis has been reported to decrease renal fibrosis in UUO models via proximal tubular cells-fibroblast crosstalk [73] as well as in myofibroblasts [25].

Recent studies have showed that WWP2 acts as a co-transcription factor involved in the regulation of SOX9 [74], RUNX2 [59], and SMAD [40], and our findings show PGC-1α as a novel target of WWP2 in myofibroblasts. We showed the specific binding of WWP2 to regulatory regions of the PGC-1α gene, whose transcription is different between WT and WWP2^-/-^ myofibroblasts. PGC-1α is a master regulator of mitochondrial biogenesis [38] and plays a crucial role in coordinating the expression of genes involved in mitochondrial function [75]. PGC-1α promotes tissue remodeling to a state that is metabolically more oxidative and less glycolytic [76], and supports tubular mitochondrial function, mitigating fibrosis progression [77]. Our data show that PGC-1α in myofibroblasts stimulates the mitochondrial respiratory function and promotes cell proliferation. These PGC-1α-mediated processes were simultaneously upregulated in WWP2^-/-^ myofibroblasts when compared with WT cells. PGC-1α levels also exhibit a negative correlation with collagen synthesis, which was also downregulated by WWP2 deficiency. While WWP2 [74] and PGC-1α [78] have been both identified as Sox9-associated proteins during chondrogenesis, the negative regulatory relationship between WWP2 and PGC-1α in fibrotic myofibroblast was not known. Furthermore, our data suggest that during fibrosis, WWP2 suppresses PGC-1α expression, as well as its downstream signaling pathways in myofibroblasts.

Using previously tested small molecules modulating PGC-1α activity, we further showed that PGC-1α mediates, at least in part, the observed effects of WWP2 on the myofibroblast polarization. PGC-1α activator ZLN005 [79] improved mitochondrial function and moderately suppressed glycolysis in renal myofibroblasts. ZLN005 also enhanced the proliferation of WT myofibroblasts while suppressing the synthesis of fibrous proteins in the presence of TGFβ1 stimulation – a result that confirms myofibroblast polarization between ‘proliferative’ and ‘pro-fibrotic’ phenotypes. Conversely, the PGC-1α inhibitor SR18292 [80] counteracted the regulatory effects of WWP2 deficiency on myofibroblasts proliferation and fibrotic activation (Figure 6N).

A few limitations in this study are pointed out here. First, while we have provided mechanistic insights into how WWP2 regulates myofibroblasts phenotypes in the fibrotic kidney, we did not characterize the transition and differentiation process of myofibroblasts during the course of CKD. Early in the disease process, the collagen matrix can undergo proteolysis, a tissue repair mechanism that can potentially reverse ongoing fibrosis. As renal fibrosis advances, the matrix proteins in fibrotic kidneys undergo modifications, becoming stiff and resistant to proteolysis [81, 82]. Our analysis focused on the late stage of renal fibrosis, and further studies are required to uncover context-specific metabolic changes in myofibroblasts at different stages of fibrosis. Second, we identified WWP2 as a transcriptional regulator of PGC-1α expression, and show how PGC-1α-target genes, identified based on gene co-expression in myofibroblasts, are under the regulation of WWP2. However, this analysis is not comprehensive. The transcription factors and co-factors mediating the regulation of PGC-1α by WWP2 are likely to be manifold, and they remain to be fully elucidated, [83]. Third, although we showed major metabolic fluxes that are affected by WWP2 (i.e. fatty acid oxidation), the primary targets of WWP2 via PGC1alpha that lead to the metabolic reprogramming of the myofibroblast remain to be identified. The metabolic changes mediated by PGC-1α in TCA cycle, and fatty acid oxidation can have broad consequences on myofibroblasts phenotype [84]. Thus, a future combination of lineage-tracing studies, a cell-type specific genomics approach, and cell subtype-specific targeting strategies will be needed to render the broad significance of myofibroblast metabolism in renal fibrosis. Finally, and most importantly, the specific pathway(s) linking cell proliferation to pro-fibrotic activation in renal myofibroblasts, remain to be discovered. Together with others, we propose that mitochondrial function has an important role in regulating these two processes that are crucial in the deposition of ECM by the myofibroblast. Future studies are required to dissect the chronology of these cellular events and identify regulators other than WWP2 and PGC-1α that contribute to myofibroblast activation and CKD progression.

In summary, these studies provide new insights into WWP2 function in renal fibrosis and its role in CKD progression. Our single cell and molecular analyses in WWP2-deficient mice provide evidence for the functional role of WWP2 in myofibroblasts and tissue fibrosis, and we propose a link between WWP2 and PGC-1α-mediated metabolic alterations in kidney fibrosis. Based on our *in vivo*, *in vitro* and clinical CKD patient studies, we propose that inhibiting WWP2 to regulate myofibroblasts metabolism might represent a promising therapeutic strategy for attenuating fibrosis progression in CKD. Therefore, WWP2 could be considered as a novel pharmacological target for the treatment of kidney disease.

