## Supplementary material for "WWP2 MEDIATES THE METABOLIC REPROGRAMMING OF RENAL MYOFIBROBLASTS TO PROMOTE KIDNEY FIBROSIS": Method

### METHODS

#### EXPERIMENTAL MODEL AND SUBJECT DETAILS

##### Human samples

Bari Cohort was enrolled from the Nephrology Unit of the University of Bari, Italy. The protocol was approved by the Independent Ethics Committee of the "Azienda Ospedaliero Universitaria Policlinico Consorziale di Bari" and was conducted in accordance with the Helsinki Declaration (Prot. N.4104/2013).

In Bari Cohort, we analyzed 34 patients with a histological diagnosis of focal segmental glomerulosclerosis (FSGS), 37 patients showing evidence of immunoglobulin A nephropathy (IgAN), 39 patients with membranous nephropathy (MN).

Nanjing cohort was enrolled from the Department of Nephrology in the Children's hospital of Nanjing Medical University in Nanjing, China. The protocol was approved by the Independent Ethics Committee of the hospital and was conducted in accordance with the Helsinki Declaration (No. 202304069-1). In Nanjing Cohort, we analysed 23 patients diagnosed with IgAN.

All patients from both cohort in the study previously gave written informed consent for the use of biopsy material for research purposes. Detailed information on those patients is provided in Supplementary Table 1. Biopsy specimens were evaluated through light, immunofluorescence and electron microscopy independently by two renal pathologists, blinded to patients' features to diagnose the disease.

##### Mouse models

Mice were bred and maintained in animal facility of Duke-NUS Medical School prior to use. Protocol with IACUC number 2021/SHS/1653 was approved by Institutional Animal Care and Use Committee of National University of Singapore, Duke-NUS Medical School, Singapore. All mice were housed in a specific pathogen–free (SPF) environment and complied with all relevant ethical regulations according to guidelines issued by the National Advisory Committee on Laboratory Animal Research. The housing room was set to a 12 hrs light/dark cycle with lights off at 8 a.m., a temperature of about 22°C and a relative air humidity of about 50%. Steps were performed to minimize animal suffering according to guidelines of the SingHealth Council on Animal Care.

With respect to the unilateral ureteral obstruction (UUO) model, mice underwent ligation of the left ureter and were sacrificed on day 7 and day 14. For the folic acid (FA) model, mice were injected with FA (250 mg/kg once, dissolved in 300 mM NaHCO3) intraperitoneally and sacrificed on day 21.

##### $WWP2^{-/-}$ mouse

$WWP2^{-/-}$ mouse line was previously generated in our lab as described in [1], and littermates were used from in-house mating in vivarium at Duke-NUS Medical School, Singapore. The background of mice is C57BL/6J (B6J). Pairs of female and male heterozygous $WWP2^{+/-}$ mice were housed together for breeding. Homozygous $WWP2^{-/-}$ and WT litters were weaned at around three weeks of age and housed in same-sex groups of four to five animals per cage. Mice were randomly assigned to experimental groups and control groups.

##### $WWP2^{Tg}$ mouse

Transgenic WWP2 mice ($WWP2^{Tg}$) overexpressed mouse WWP2 full-length isoform under the immediate early promotor of cytomegalovirus (PCMV IE) on the pIRES2-EGFP Vector. In detail, 2 ml plasmid saline solution was injected in mice via tail vein injection at the dosage of 4 mg/kg within 2 seconds. Mice of 8 weeks were randomly assigned to $Control^{Tg}$ and $WWP2^{Tg}$ undergoing plasmid injection containing control and WWP2 DNA insert, respectively. The efficiency of WWP2 overexpression in the kidneys was examined 24 hrs post injection by randomly selecting 3 mice from each group; the UUO modelling was established 24 hrs post injection in the rest of the mice.

##### Primary myofibroblasts culture

Murine renal (myo)fibroblasts were derived from both kidneys of mice. Minced kidney pieces (1-3mm$^3$) were placed in 6 cm dishes with DMEM supplemented with 20% fetal bovine serum for less than 10 days to generate mice renal (myo)fibroblasts (P0) and passaged to P1 and P2 in DMEM supplemented with 10% fetal bovine serum. In each experiment, all the cells from $WWP2^{-/-}$ and WT kidneys were cultured at the same time using the same procedure (as described above). Since fibroblast-to-myofibroblast conversion occurs with each cell passage using common cell culture method [2], our primary isolated myofibroblasts and cultured cells presented myofibroblasts features, and thus we refer to them as myofibroblasts.

P2 renal myofibroblasts were used for experiments at ~80% confluence. $WWP2^{OE}$ cells were generated with infection of lentivirus which carried anti-puromycin gene and WWP2 full-length isoform DNA, and the lentivirus with scramble DNA as control. Seventy-two hours post infection, 4 μg/ml puromycin was added in the medium to deplete the non-infected cells. To activate the myofibroblasts to myofibroblasts, cells were treated with transforming growth factor-β1 (TGFβ1) human at a concentration of 5ng/μl for 24 or 72 hours as indicated. To inhibit or activate PGC-1α function, cells were treated with 10-20uM ZLN005 and SR18292 at 2.5-10uM for 30min prior to TGFβ1 treatment for 72h, respectively.

#### METHODS DETAILS

##### Renal histopathology

Masson's trichrome, Sirius Red were performed on paraffin-embedded sections. Images were acquired by the Aperio ScanScope CS2 device (Aperio Technologies, Vista, CA, USA), and digital slides were analyzed with ImageScope V12.1.0.5029 (Aperio Technologies). We excluded for the analysis peri-vascular regions, Bowman's capsule and the limits of each biopsy section by drawing tools. The positive area was expressed as the ratio of number of positive (NP) area over the total area analyzed (NP/Area).

##### Hydroxyproline Assay

The amount of total collagen in the kidneys was quantified using the Quickzyme Total Collagen assay kit (Quickzyme Biosciences). The assays were performed according to the manufacturer's protocol. The levels of HPA collagen were normalized the weight of kidneys tested and expressed as μg/mg.

##### Immunochemistry and immunofluorescence staining

For the immunohistochemical evaluation of WWP2 in human kidneys, 4 μm-thick paraffin-embedded renal tissue underwent deparaffination and heat mediated antigen retrieval (EDTA 1mM, pH=8.00). After epitope unmasking, the slides were incubated with $H_2O_2$ (0.3%) and then with Triton X (0.25%), protein block solution (Dako), and the primary antibody (1:50 dilution in PBS). Primary antibodies were detected by the Peroxidase/DAB Dako Real EnVision Detection System, according to the manufacturer's instructions (Dako). Renal sections were counterstained with Mayer hematoxylin and mounted with glycerol (Dako Cytomation, Glostrup, Denmark). Negative controls were prepared using isotype control antibodies.

For the immunofluorescence staining for kidney samples, sections of 5 um thickness of kidneys were fixed with 10% Neutral Buffered Formalin and processed using Leica automatic tissue processor. Following dewaxing and rehydration, sections were heated in citrate buffer for antigen retrieval for further antibody incubation. For cell sections, P2 renal fibroblasts were seeded onto 8 well removable chamber slides (#80841, ibidi) and fixed with ice cold acetone for 30min at room temperature. Antibodies used for immunofluorescent staining are as follows: anti-ACTA2 (Sigma-Aldrich, 1:100), anti-WWP2 (Bethyl Laboratories, 1:100), anti-Vimentin (Abcam, 1:100). Images were visualised using Secondary Antibodies (A11008, A11004 and A10525; Thermo Fisher Scientific) at 1:500 for 2h at room temperature. Sections were imaged under a Leica Fluorescent microscope.

##### Oil-red O staining

Renal myofibroblasts were seeded onto 24 well plate at a density of 8000 cells/well. After treatment with TGFβ1 (5ng/μl) for 72 h, cells were fixed with 4% PFA for 30 min at room temperature. After brief rinsing with PBS, plates were incubated with Oil Red O solution (0.5% in isopropanol, #O1391, Sigma- Aldrich) for 20 min at room temperature. Images were obtained using a light microscope after rinsing the plates with $dH_2O$. Quantification of Oil Red O stain extracted from stained cells was carried out by eluting the stain from cells in 1 mL of 100% isopropyl alcohol, and then measuring the absorbance of the stain against a blank (100% isopropyl alcohol) at 500 nm.

##### qRT-PCR analysis

Total RNA was extracted from snap- frozen tissue and primary cells using the RNeasy mini kit (Qiagen), and cDNA was prepared using iScript cDNA synthesis kit (primer specific, BIORAD) according to the manufacturer's instructions. Fast SYBR-Green master mix (BIORAD) was used for the analysis of gene expression using the BIORAD CFX RT-PCR system. The primers used in the experiment are listed in the Resource Table. 18S was used to normalize the relative gene expression, and $2^{-\Delta\Delta Ct}$ method was used to measure the fold change.

##### Western blot analysis

Cell lysates were obtained from kidney tissues and cells using RIPA buffer (Thermo Fisher Scientific) supplemented with protease (Sigma-Aldrich) and phosphatase inhibitor cocktails (ROCHE). Lysates were subjected to 4-12% SDS-PAGE electrophoresis after Bradford quantification. Blotting of the membrane was performed using anti-WWP2 (Bethyl Laboratories, 1:500), anti-ACTA2 (Sigma-Aldrich, 1:10,000), anti-Vimentin (Abcam, 1:500), anti-Periostin (Novus Bio, 1:500), anti-Fibronectin (Sigma-Aldrich, 1:500), anti-Col1a1 (SouthernBiotech), anti-PGC1a (Abcam), anti-CyclinA (GeneTex), anti p21(GeneTex) after transfer onto a nitrocellulose membrane at 100 V for 1 hr. Blots were visualized with anti-Rabbit HRP (Merck, 1:5000) and anti-Mouse HRP (Merck, 1:5000) on a Kodak automated developer and ChemiDoc MP imaging system (Bio-RAD) after blocking with 5% non-fat dry milk for 1 hr at room temperature using Pierce ECL Chemiluminescent substrate (Thermo Fisher Scientific) and Immobilon Forte Western HRP substrate (Merck), and then quantified using densitometry using ImageJ software. Anti-Tubulin (Sigma-Aldrich, 1:5000) and anti-GAPDH (Abcam, 1:5000) were used as loading controls. Full and unprocessed scanned images of the blots are provided in the Source Data file.

##### Flow cytometry analysis

Mouse kidneys were perfused, excised, minced, and digested with Collagenase II (Worthington Biochemical Corporation) and Dispase II (Roche). Tissue mixture was mechanically disrupted and filtered through 70 µm cell strainer to obtain single cell suspension. Cultured myofibroblasts were seeded onto 10cm dishes, and were harvested for single cell suspension after treatment with TGFβ1 (5ng/µl) for 72h.

After blocking with CD16/32 (Thermo fisher, 1:100) at RT for 10 min, cells were fixed and permeabilizated using Intracellular Fixation & Permeabilization Buffer Set (eBioscience). Cells were collected by centrifugation, subjected to live-death dye and antibody staining, and analysed (or sorted) by flow cytometry using BD FACS ARIA system; the data were analysed using FlowJo™ v10 software.

##### Cell proliferation assay

Cell proliferation was quantified by MTS assay (Promega) according to the manufacturers' protocol. For cell cycle assay, cells were harvested and fixed using the Propidium Iodide Flow cytometry Kit (Abcam) and measured using BD FACS ARIA system. Data were analysed using the ModFit LT software (DNA Modelling System).

##### ChIP-seq and ChIP-qPCR analyses

Chromatin immuno-precipitation (ChIP) assay was performed using Magnetic ChIP Kit (Pierce) with anti-WWP2 antibody (#A302-936A, Bethyl Laboratories). Briefly, P2 WT renal fibroblasts were grown for 80-90% confluence. After treatment with TGFβ1 (5ng/µl) for 24 h, cells were harvested by trypsinisation. DNA and proteins in the cells (~$10^6$ each sample) were cross-linked using 1% (v/v) formaldehyde and then sonicated in lysis buffer to obtain 200 bp–500 bp long DNA fragments. Supernatant was incubated with control IgG or WWP2 antibody beads at 4°C, or kept as input references. Reverse-crosslinking and purification of DNA was performed.

ChIP-seq libraries were constructed and sequenced using Illumina HiSeq 4000 sequencer, resulting in paired-end FASTQ files containing sequence reads of 150bp length. These FASTQ files were mapped to mouse reference genome GRCm38 using STAR aligner. PRC duplicated reads were removed by Picard MarkDuplicates function. Peak calling was performed by MACS2 callpeak function[3] using the following parameters: "-qvalue 0.001", "--keep-dup auto", and "--call-summits". The genomic location of the peak maximum was extended 250pb both upstream and downstream, defining a 501 bp long sequence centered on each ChIP-seq peak. The transcription factor binding site (TFBS) motifs discovery and annotation was analyzed by the RSAT peak-motif online server [4], Regulatory Sequence

Analysis Tools (RSAT)[5] using default parameters, and JASPAR core nonredundant vertebrates and ENCODEdatabases[6]. The results of the TFBS motifs identified are listed in Supplementary table 2.

##### Bulk RNA-seq data generation and analysis

Primary renal myofibroblasts (P2) were harvested and RNA was extracted using the RNeasy mini kit (Qiagen) following the manufacturer's instruction. Libraries were generated with the poly-A selection method (mRNA Direct kit, Life Technologies), and sequenced using the NovaSeq 6000 S4 platform (23150 bp) with a target of 30 million reads per library.

Libraries were sequenced using paired-end 150bp sequencing strategy. Reads were mapped to the mouse genome GRCm38 (mm10) v89 using STAR[7] aligner, and counts per gene were quantified by featureCounts[8] with settings of pair-end (-p) and exon mapping (-t exon). Genes were considered detected if they have at least 5 counts in at least 5 samples. Counts were then normalized for sequencing depth and RNA composition across all samples by DESeq2.

Differential gene expression analysis was performed between two sample groups (wildtype (WT) and WWP2$^{-/-}$ mouse) by Wald test, the default test is DSeq2. After multiple testing correction, genes with a false discovery rate (FDR) <0.05 were considered significantly differentially expressed unless otherwise indicated. Functional enrichment analysis test of the differentially expressed genes in fibroblast cells between WT and WWP2$^{-/-}$ was performed by clusterProfiler, and visualisation of enrichment results was performed by gseaplot function. Pathways relevant to ECM and cell proliferation were selected from the MSigDB curated genesets[9; 10].

##### Metabolic function analysis

Metabolic measurements of myofibroblasts were obtained by real-time oxygen consumption rate (OCR) and extracellular acidification rate (ECAR) using a Seahorse XFe96 Analyzer (Agilent). Briefly, 10,000-20,000 cells were seeded into each well of a XF96 cell culture microplate (102416-100, Agilent). For Cell Mito Stress Test analysis, the culture medium was switched to Seahorse XF DMEM assay medium with 1 mM pyruvate, 2 mM glutamine and 10 mM glucose, and then placed into a $CO_2$ free 37°C incubator for 1 h prior to the assay. The OCR measurements were recorded using serial injections of Oligomycin (10 µm), FCCP (1 µM) and antimycin A (0.5 µM) plus rotenone (0.5 µM). For the Glycolysis Stress Test analysis, the culture medium was changed to Seahorse XF DMEM supplemented with 2 mM glutamine, and then placed into a $CO_2$ free 37°C incubator for 1 h prior to the assay. The ECAR measurements were recorded using serial injections of Glucose (15 mM), Oligomycin (10 µm), 2-DG (50 mM). Upon completion of each Seahorse XF assay, the medium was discarded, and the cells were incubated in 0.5mg/ml MTT for 3 hrs for normalization. The number of cells per well was normalized by MTT assay, which was used to measure cellular metabolic activity.

##### Metabolomics analysis

After treatment with TGFβ1 (5ng/µl) for 72 h, primary renal myofibroblasts (P2) were harvested in 0.6% formic acid and sonicated with 10 cycles (30 sec ON, 30 sec OFF). Acetonitrile was added in a 1:1 ratio to the formic acid volume, and the mixture was vortexed for homogeneity. We used a metabolomics approach to quantify acetylcarnitines, amino acids, glucose molecules and organic acids involved in fatty acid oxidation, amino acid metabolites, glycolysis, and TCA cycle intermediates, which was carried out in the Metabolomics Facility at Duke-NUS Medical School.

The glycolytic intermediates were separated by capillary ion chromatography (IC) on a Dionex ICS-4000 capillary system (Thermo Fisher Scientific, USA) and monitored on a Thermo Q Exactive Plus Q-Orbitrap HRMS (Thermo Fisher Scientific, USA). For amino acids and acylcarnitine panel, the samples were run with a C18 column (Phenomenex, 100 x 2.1 mm, 1.6 µm, Luna® Omega) on 1290 Infinity LC system (Agilent Technologies, USA) coupled with quadrupole-ion trap mass spectrometer (QTRAP 5500, AB Sciex, USA). Trimethylsilyl derivatives of organic acids were separated by gas chromatography (Agilent Technologies, HP 7890A) and quantified by selected ion monitoring on a 5975C mass spectrometer using stable isotope dilution. Data acquisition and analysis were performed on an MassHunter Workstation B.06.00 (Agilent), MultiQuant™ 3.0.3 software (AB Sciex, DC, USA) and using the TraceFinder software (Thermo Fisher Scientific, USA). Data were normalized with respect to the protein concentration. Statistical differences between WT and WWP2$^{-/-}$ groups were assessed by two-tailed non-parametric Mann-Whitney U test, and P-values were corrected using the Benjamini-Hochberg-correction.

#### Single-cell RNA-seq data generation and analysis

*Library generation and sequencing.* Mouse kidneys were perfused, excised, minced, and digested to obtain single cell suspensions as described above. Cells were collected by centrifugation, subjected to live-death dye, and sorted by flow cytometry using BD FACS ARIA system. Isolated living single cell suspensions were converted to barcoded scRNA-seq libraries by using the Chromium Single Cell 3′ Library, Gel Bead & Multiplex Kit, and Chip Kit V3, loading an estimated 7,000–12,000 cells per library/well, and following the manufacturer's instructions. Indexed libraries were sequenced using Illumina NovaSeq 6000 platform, where 150bp pair-end sequences were obtained. Sequencing reads were aligned and quantified to the mouse genome GRCm38 (mm10-3.0.0 provided by 10x Genomics) using 10x Genomics Cell Ranger count.

*Data pre-processing and cell annotation.* Quality control and pre-processing was performed individually for each of the four samples (WWP2$^{-/-}$, n=2 and wildtype (WT), n=2). Genes present in less than 50 cells and cells below the 10$^{th}$ and above the 90$^{th}$ percentile in terms of both the number of genes and number of counts were filtered out. Cells with percentage of mitochondrial content greater than 40% and haemoglobin gene accounting more than 1% of total counts were also filtered out. In addition, cells above the 99$^{th}$ percentile in terms of percentage of total counts being stress genes[11] were also removed. Seurat (v4.0) was used to normalize the filtered data using normalization method of “LogNormalize” and scale factor of 10,000. The top 5,000 highly variable genes were used for the principal component analysis (PCA), and top 50 principal components were used for Uniform Manifold Approximation and Projection (UMAP) dimensionality reduction. Doublet identification and the calculation of doublet score were performed using co-expression based doublet scoring (cxds)[12]. Clusters with median doublet score of greater than the 60$^{th}$ percentile were denoted as potential doublet clusters and filtered out. Small outlier clusters with fewer than 1% of total number of cells were also filtered out. For cell type annotation, markers from a published scRNA-seq study of murine kidney using the UUO model[13] were used as reference. Cell type scores were calculated using Seurat (v4.0) *AddModuleScore()* function with random seed set at 42 and control gene set size of 500. Final cell type annotations were obtained by identifying the cell type with the highest normalized score (dividing each cell type score by the maximum cell type score). The cell type annotations were further confirmed by using two other scRNA-seq datasets from murine kidney [14; 15]. Renormalization of the filtered data was performed using Seurat (v4.0) with the clustering resolution of 0.8. We recalculated doublet score using the cxds approach as above. Clusters with median doublet score of greater than the 95$^{th}$ percentile were denoted as potential doublet clusters and filtered out. This yielded 4 processed data matrices corresponding to initial the 4 scRNA-seq 10x libraries. The 4 processed datasets were merged, and the same Seurat pipeline highlighted above was implemented using a clustering resolution of 1.0. After merging, mixed clusters were filtered out. In addition, batch-specific clusters, which have 1 batch dominating at least 75% of cells, were removed. This process was done recursively until there were no batch-specific or doublet clusters present. This merged dataset was renormalized using the same Seurat (v4.0) pipeline with a clustering resolution of 1.0. Lastly, small clusters (accounting for fewer than 0.25% of total cell number) and 12 outlier cells were filtered out, giving us a final processed dataset consisting of 17,058 genes and 74,585 cells.

*Differential expression analysis and functional enrichment tests.* Differential expression between WWP2$^{-/-}$ and WT was performed by running Seurat’s *FindMarkers()* function with minimum log fold change of 0 and minimum percentage set to 0. Functional enrichment test in the differentially expressed genes in fibroblast between WT and WWP2$^{-/-}$ was performed by *clusterProfiler* using pathways in the Reactome database[16] (release 83). Enriched terms were grouped by *treeplot()* function that hierarchically clustered terms based on the pairwise similarities, which were calculated by the *pairwise_termsim()* function. The final plot of hierarchical clustering of pathways was further manually curated by merging similar pathways/processes terms based on the similarity of enriched genes. Full enrichment results are reported in Supplementary Table 3.

*External datasets of human kidney fibroblasts.* Two human single cell kidney datasets[17; 18], including fibroblasts from both CKD and healthy patients kidneys were used in our study, and the original cell annotation provided by authors was used for downstream analysis. For the dataset “human_CD10negative_final.RData” (https://zenodo.org/record/4059315), we used fibroblasts annotated as "Fibroblast" and "Myofibroblast" cells in “Annotation.Level.3”. For the Kidney Precision Medicine Project (KPMP) dataset “snRNA_kidney_atlas_KPMP.rds” (https://cellxgene.cziscience.com/collections/bcb61471-2a44-4d00-a0af-ff085512674c), we used cells annotated as "kidney interstitial fibroblast" in “celltype”. For both datasets, data from patients (both healthy and CKD) with less than 20 fibroblasts were removed in downstream analysis.

*Pseudo-time analysis.* Slingshot was used for pseudo-time analysis of cell state changes. ECM gene sets (NABA_MATRISOME, NABA_CORE_MATRISOME, NABA_BASEMENT_MEMBRANE, NABA_ECM_REGULATORS, NABA_ECM_GLYCOPROTEINS, NABA_COLLAGENS, NABA_PROTEOGLYCANS)[9] were downloaded from GSEA MSigDB[19]. The ECM gene sets were integrated and the average expression scores of ECM gene sets visualized on UMAP reduction dimension along the trajectory inferred by Slingshot.

*Cell phase estimation and visualization.* The cell phase was determined for each cell using the Revelio method described in [20]. This method extracts a set of marker genes for each cell phase (M to G1, G1 to S, S, G2, G2 to M) and, for each cell, calculates an expression score for each phase. The cell status was assigned based on the maximum z-score between the phases. Using dimensionality reduction method, all cells were projected into two-dimension based on their dynamical components (DCs), which are analogue to principal components. The visualization of the “pseudo-time” of cell phases was the revolution of cartesian to polar transformation of the 2D DC plot. The line summarizing expression score in the cell cycle plots in Figure 3H-I was added by *geom_smooth()* function, which uses a generalized additive mode smoothing (GAM) method. To tested if the gene expression changes significantly along the cell cycle phases (G0>G1>S>G2>M) we employed the WAVK trend test[21] which is implemented by R package *funtimes* v9.1[22]. WAVK is a nonparametric test to detect (non-)monotonic parametric trends in time series; here, the time series is represented by the pseudotime trajectory alongside the cell cycle phases (G0>G1>S>G2>M), and the null hypothesis tested the by WAVK trend test is no change in trend along the pseudotime trajectory. For illustration purpose, phases M to G1 and G1 to S are marked as G0/1, and phases G2 and G2 to M are marked as G2/M.

*Estimation of cell metabolomic activities from scRNA-seq data*. We used the COMPASS algorithm[23] to estimate the cellular metabolic states based on single-cell RNA expression and its inference of flux balance. For each cell, COMPASS calculates the activity scores of a comprehensive list of metabolic reactions. The difference of these activities scores between WWP2$^{-/-}$ and WT group were tested by non-parametric Wilcoxon rank-sum test, and the effect size was estimated by Cohen’s D. For each cell, we also calculated the correlation of WWP2 mRNA expression (normalised counts with scale factor 10k) and the metabolic activity score by Spearman correlation coefficient.

*Correlation and co-expression network modules.* We used the bigSCale analytical framework[24] to estimate the gene-gene correlation coefficient (*r*) and infer the network of genes centred on PGC-1α (*PPARGC1A*) in myofibroblasts scRNA-seq data from WT and WWP2$^{-/-}$ mouse kidneys. The genes correlated ($|r| > 0.5$, $P<0.001$) with PGC-1α defined two gene co-expression networks - positively correlated network and negatively correlated network. Each was further clustered into different sub-modules using multi-level modularity optimization algorithm, cluster_louvain (igraph package[25] with parameter resolution = 1 which yields to fewer clusters) and resulted to 2 main interrelated modules. In detail, module 1 and 2 include genes positive correlated with PGC-1α, while module 3 and 4 include genes negatively correlated with PGC-1α. For illustration purposes, genes in module 1 and 2 or module 3 and 4 were grouped together, and only the genes with gene-gene correlations with $|r| > 0.6$ were included. The genes highlighted in the Figure 6E-F are those that are more strongly correlated with PGC-1α (i.e., $|r| > 0.8$)**.** Differences in module gene expression between WWP2$^{-/-}$ and WT were estimated using bulk RNA-seq data (see Bulk RNA-seq data generation and analysis above) generated in cultured renal fibroblast, TGFb1 treated and untreated, from WT and WWP2$^{-/-}$ mice. TPM (transcript per million) expression for each gene in the module was obtained, and normalized by z-score across all samples. For each module, global expression differences between between WWP2$^{-/-}$ and WT were tested by non-parametric Wilcoxon rank-sum test on the z-score normalized data.

#### QUANTIFICATION AND STATISTICAL ANALYSIS

Data are expressed as mean ± standard deviation (SD), unless otherwise indicated. The applied statistical tests (two-tailed non-parametric Mann-Whitney U test, two-tailed non-parametric Wilcoxon rank-sum test, the chi-square test (for proportions)), which were dependent on the number of groups being compared and the study design, are indicated in each figure legend. R (v4.2.3) statistical programming language or SPSS (v26) Statistical Software Platform were used in all statistical analyses. All experiments requiring the use of animals, directly or as a source of cells, were subjected to randomization. The experimenters were blinded to the grouping information. All *in vivo* model analyses were conducted with sample sizes of at least 8, determined through power calculation analysis. In vitro experiments were independently replicated a minimum of three times, as stated in the figure legends. Statistical significance is indicated by P-value, where * denotes $P < 0.05$, and ** denotes $P < 0.01$ unless otherwise indicated.

#### RESOURCE AVAILABILITY

##### Lead contact

##### Materials availability

This study did not generate new unique reagents. All the key resources used are listed in Supplementary table 4.

##### Data and code availability

Raw (FASTQ files), pre-processed data (count matrix) and metadata of the scRNA-seq datasets, bulk RNA-seq data generated in primary myofibroblasts and Raw and processed bulk ChIP-seq data have been deposited in NCBI's Gene Expression Omnibus. GEO Series accession number is pending. Scripts for the analysis pipeline used for single-cell clustering, data visualization and generation of all major figures in this study were written in R, and code is available at Github.
