## Supplementary Figures for "WWP2 MEDIATES THE METABOLIC REPROGRAMMING OF RENAL MYOFIBROBLASTS TO PROMOTE KIDNEY FIBROSIS"

### Supplementary Figure 1

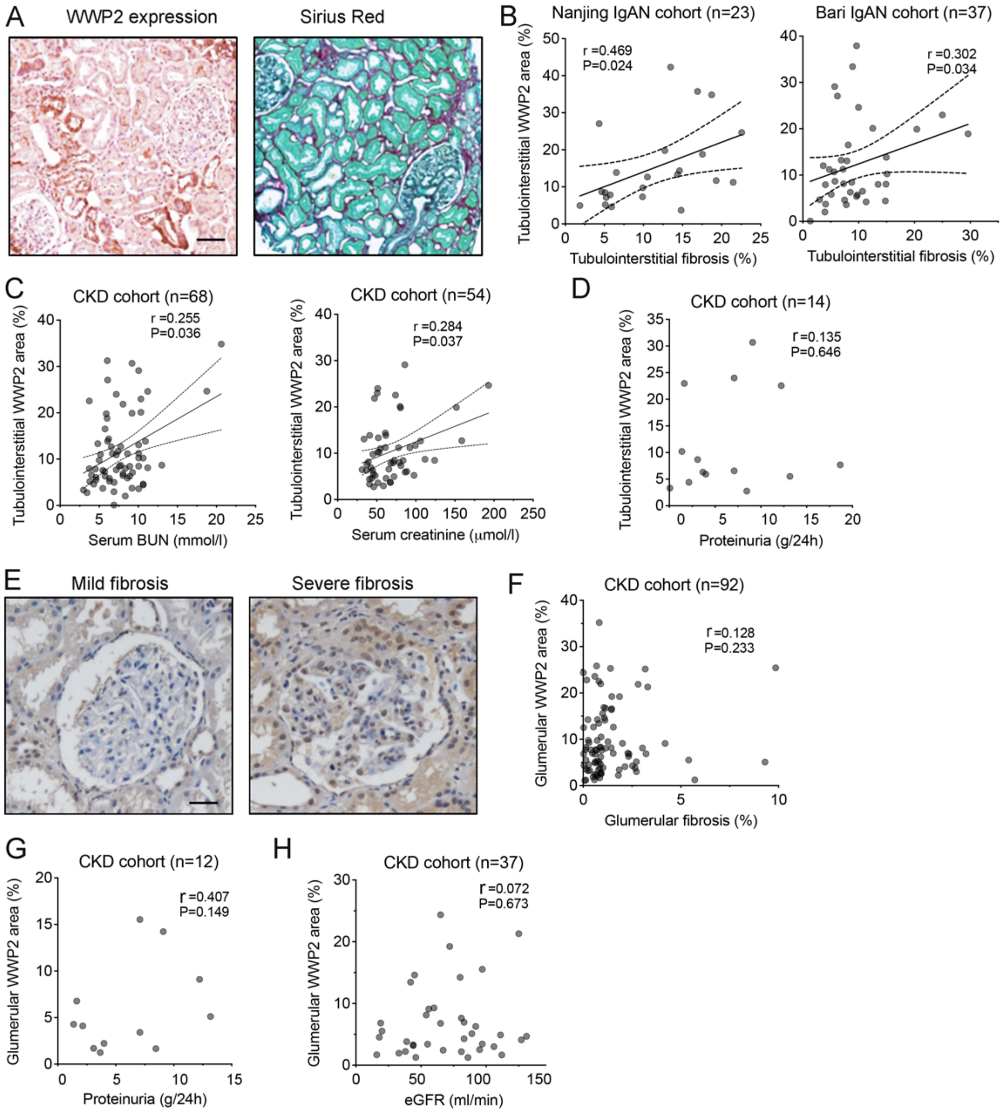


**Supplementary Figure 1. Expression of WWP2 in the kidney of patients with chronic kidney disease (CKD) and UUO-mice (related to Figure 1)**

(A) Representative images of WWP2 immunostaining (*left*) and Sirius Red staining (*right*) in human kidney biopsy samples, utilized for quantitative analysis for WWP2-positive area and fibrosis-positive area. Scale bars, 25 μm.

(B) Positive correlation between tubulointerstitial WWP2-positive area and fibrosis-positive area in the Nanjing IgAN cohort (n=23, *P*=0.024, *r*=0.469; *left*) and the Bari IgAN cohort (n=37, *P*=0.034, *r*=0.302; *right*), which was determined by immunostaining and Sirius Red staining in Supplementary Figure 1A using ImageScope. Data points represent measurements of individual section obtained from each patient with CKD.

(C) Positive correlation of tubulointerstitial WWP2-positive area with serum BUN levels (n=68, *P*=0.036, *r*=0.255; *left*) and with serum creatinine levels (n=54, *P*=0.037, *r*=0.284; right) in patients with CKD (multi-centre cohort).

(D) Correlation of tubulointerstitial WWP2-positive area with proteinuria levels in a subgroup of patients with CKD (n=14, *P*=0.646, *r*=0.135).

(E) Representative immunostaining images of WWP2 in glomerular area from human renal biopsy samples, illustrating mild and severe glomerular fibrosis, respectively (110 patients with CKD recorded from Bari cohort). Scale bars, 25 μm.

(F-H) Correlation of glomerular WWP2-positive area with glomerular fibrosis-positive area (n=92, *P*=0.233, *r*=0.128; panel F), proteinuria levels (n=12, *P*=0.233, *r*=0.128; panel G) and eGFR levels (n=37, *P*=0.673, *r*=0.072; panel H) in patients with CKD from Bari cohort.

### Supplementary Figure 2

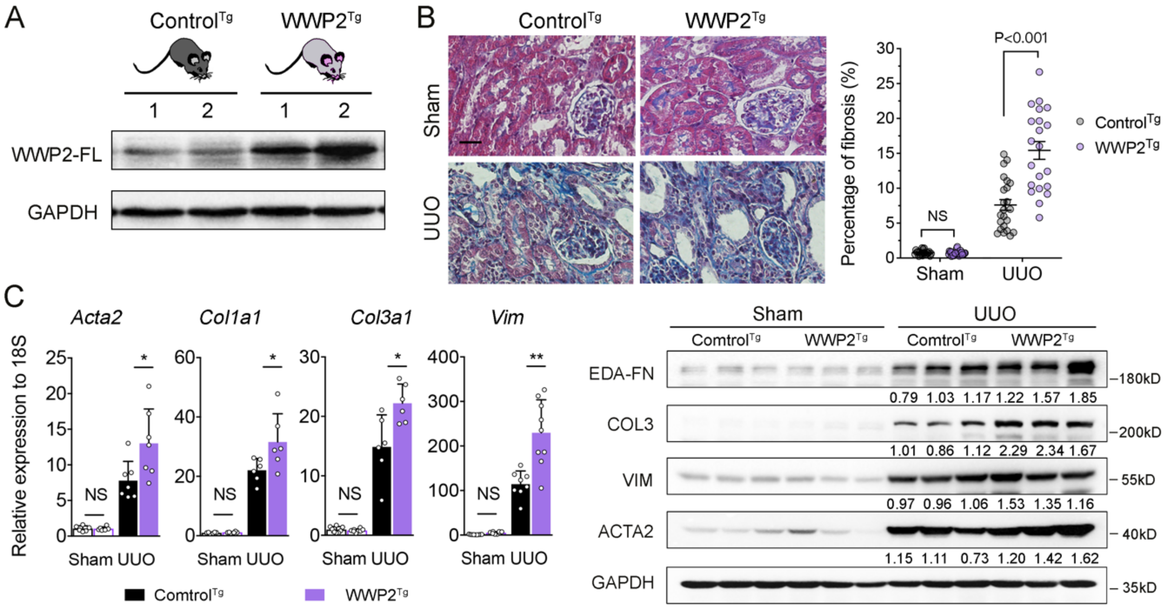


**Supplementary Figure 2. WWP2 overexpression exacerbates renal fibrosis *in vivo* (related to Figure 1)**

(A) Representative western blot showing the levels of WWP2 in kidney tissue from Control$^{Tg}$ and WWP2$^{Tg}$ mice. Control$^{Tg}$ and WWP2$^{Tg}$ refer to mice injected with plasmids containing control and WWP2 DNA, respectively (see Methods for details).

(B) Representative images of Masson's trichrome staining in kidney tissue sections from Sham and UUO-treated mice (10 days), with WT and WWP2$^{-/-}$ genotypes (*left*, scale bar, 30 µm), followed by quantitative analysis (*right*). Each dot indicates the average fibrosis-positive area from the section per mouse, and five non-overlapping fields were taken. Values are reported as mean ± SEM. P-values calculated by two-tailed Mann-Whitney U test.

(C) The expression of ECM genes in kidney tissues from UUO and control mice (n=6-9, each group). *Left*, expression changes were determined with RT-qPCR; *right*, representative western blot for protein levels analysis. The values are reported as mean ± SD. P-values calculated by two-tailed Mann-Whitney U test.

### Supplementary Figure 3

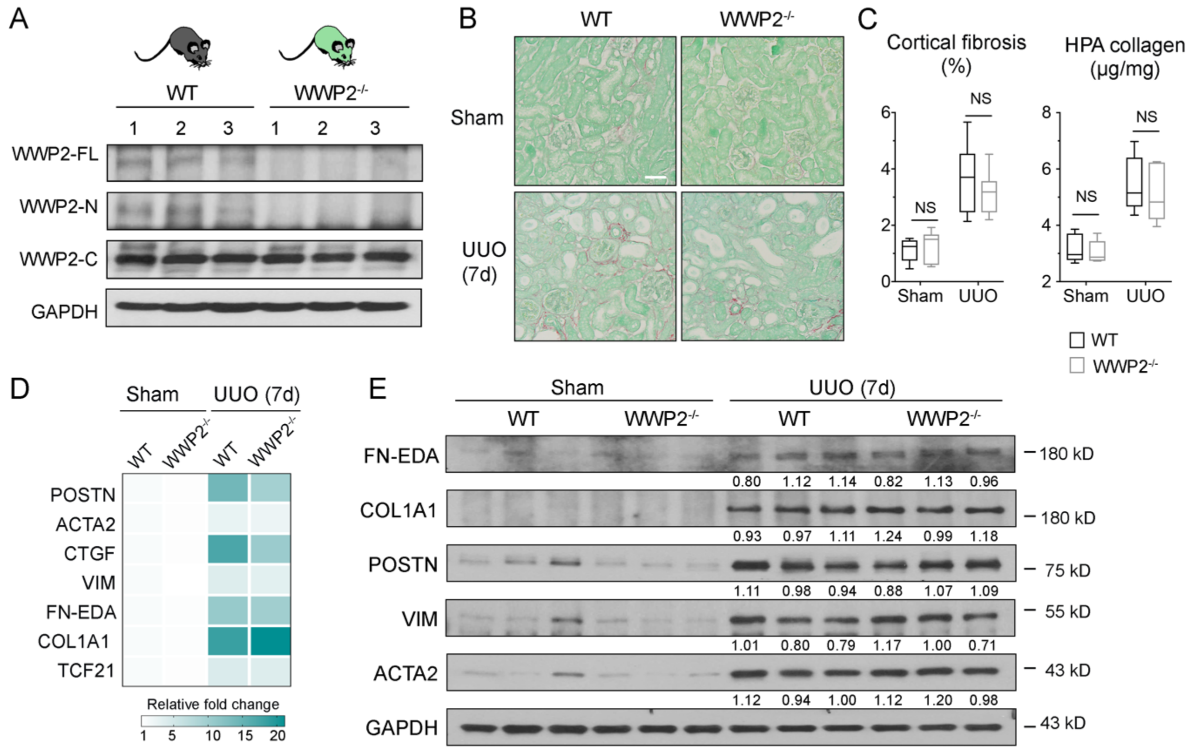


#### Supplementary Figure 3. Effect of WWP2 deficiency on renal fibrosis at 7 days of UUO (related to Figure 2)

(A) Representative western blot showing the levels of WWP2 in kidneys from WT and WWP2$^{-/-}$ mice. WWP2$^{-/-}$ kidneys preserve WWP2 C-terminal isoform, while lack the WWP2-full length (FL) and -N isoforms.

(B) Representative Sirius Red images of WT and WWP2$^{-/-}$ mouse kidneys following UUO model for 7 days. (n=6 images recorded for each condition). Scale bars, 10 µm.

(C) Quantitative analysis of cortical fibrosis-positive area (left, %) and HPA collage levels (right, µg/mg) WT and WWP2$^{-/-}$ mouse kidneys following UUO model for 7 days. The data values are summarized using box-and-whisker plots (n=6). P-values were calculated by two-tailed Mann-Whitney U test. NS, *P*>0.05.

(D) Heatmap showing of mRNA expression of ECM genes, determined with RT-qPCR, in kidney from WT and WWP2$^{-/-}$ mouse following UUO model for 7 days. For each gene, relative fold change was calculated with respect to Sham WT and averaged across biological replicates (n=6, each group). No statistically significant differences (P<0.05) were detected between WT and WWP2$^{-/-}$ groups.

(E) Representative western blot for ECM proteins in UUO kidneys from WT and WWP2$^{-/-}$ mouse kidneys following UUO model for 7 days. No statistically significant differences (P<0.05) were detected between WT and WWP2$^{-/-}$ groups.

### Supplementary Figure 4

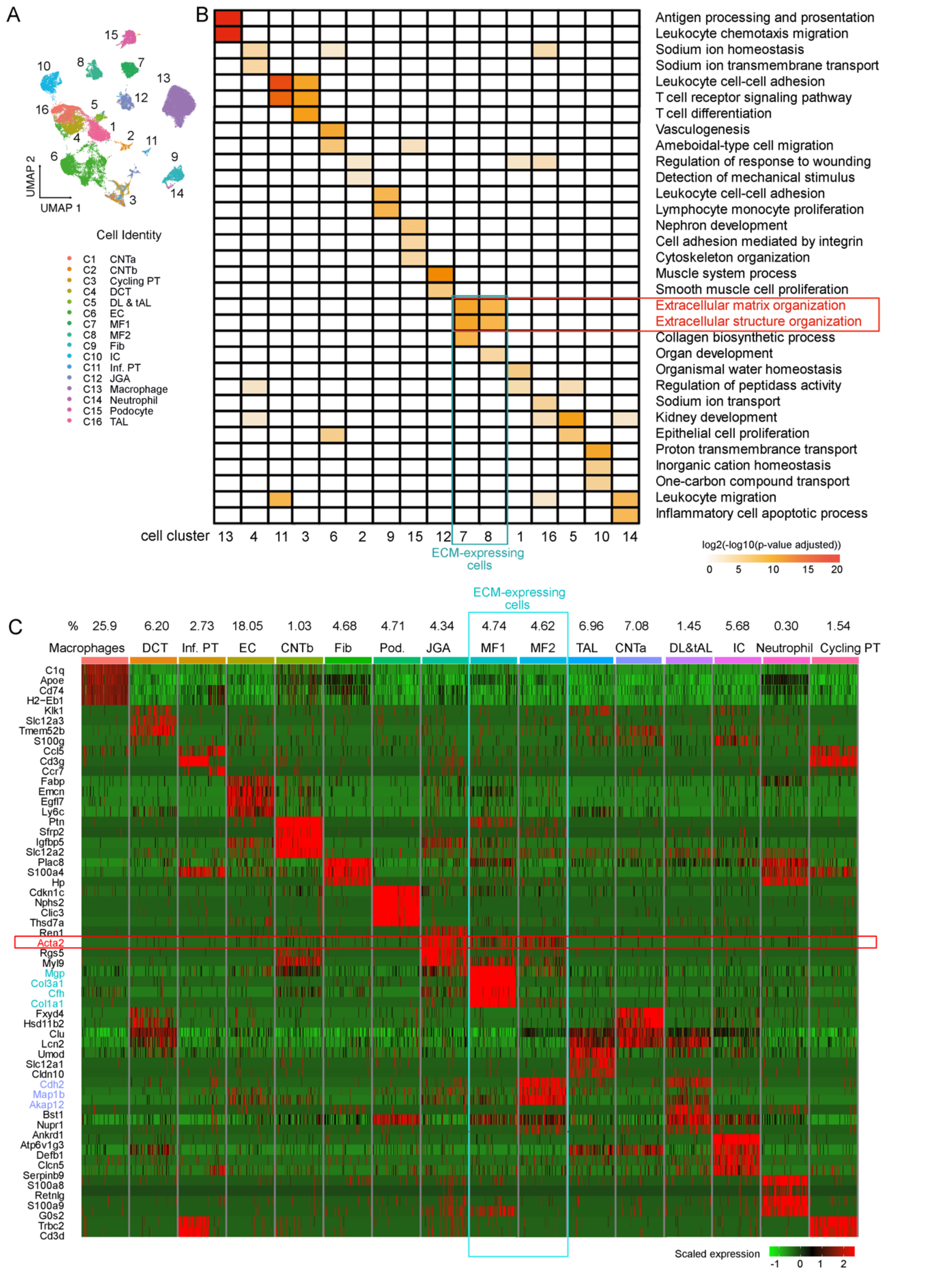


**Supplementary Figure 4. Single-cell transcriptomics landscape of kidney cells derived from UUO-mice (related to Figure 3)**

(A) Uniform Manifold Approximation and Projection (UMAP) representation of 74,585 kidney cells profiled in WT (n=2) and WWP2$^{-/-}$ (n=2) UUO-mice, showing 16 major cell types. CNT:

connecting tubule; PT: proximal tubule; DCT: distal convoluted tubule; DL & tAL: descending limb and thin ascending limb; EC: endothelial cells; MF: myofibroblasts; Fib: fibroblasts; IC: intercalated cells; Inf. PT: inflammation proximal tubule; JGA: juxtaglomerular cells; TAL: thick ascending limb. Top 4 marker genes are reported in panel C.

(B) Heatmap of enriched pathways identified in each renal cell-type cluster. Only the top 2 significant terms are shown, and the significance of enrichment is represented by $\log_2(-\log_{10}$(adjusted p-value)). Pathways involved in ECM process are highlighted (red box) and were identified in the ECM-expressing clusters (cluster 7 and 8, light blue box).

(C) Heatmap of the scaled gene expression of top 4 marker genes for each cell cluster (panel A), based on average $\log_2$ fold change (FC) in gene expression. The marker gene ACTA-2 is highlighted (red box), which is detected in JGA cluster and in the ECM-expressing cluster 7 and 8 (light blue box).

**Supplementary Figure 5**

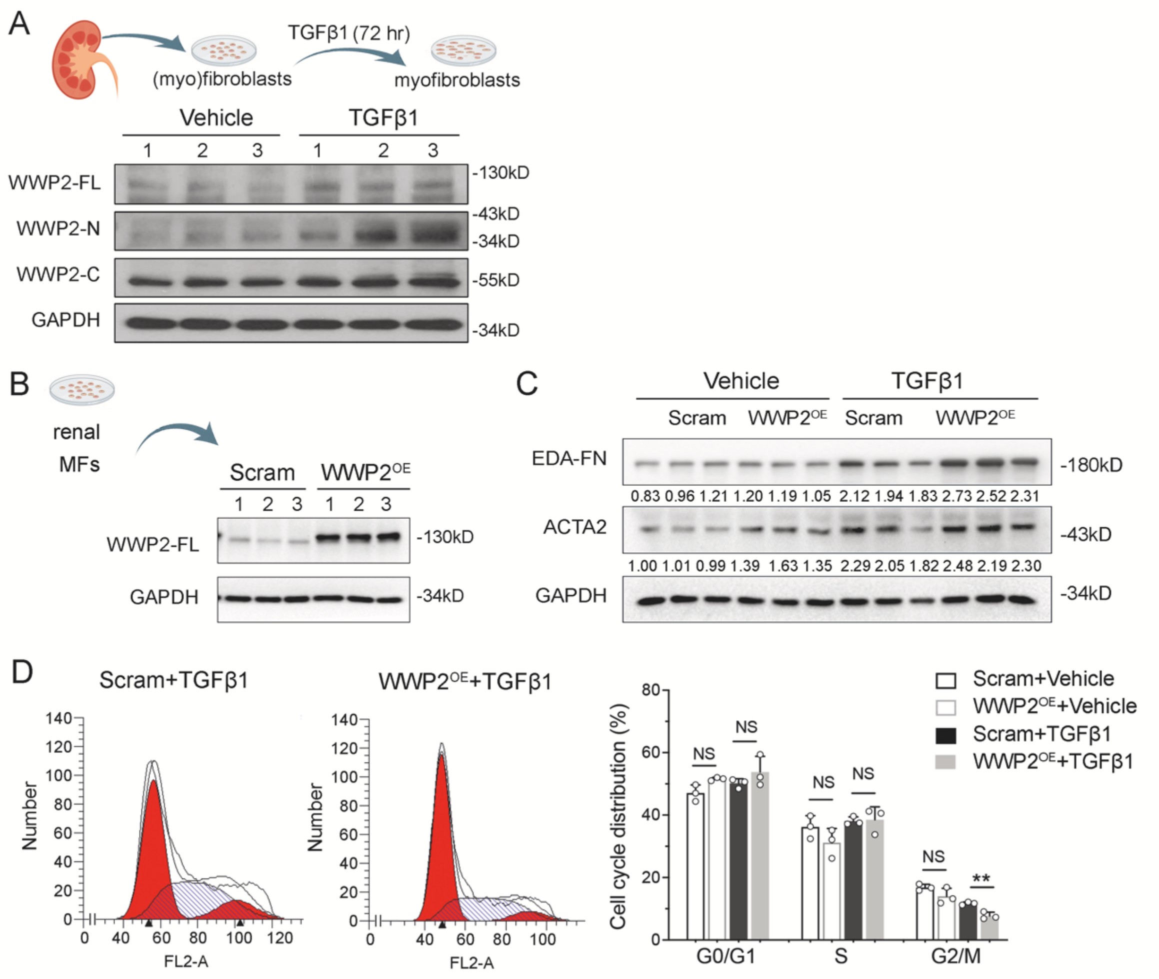


**Supplementary Figure 5. WWP2 overexpression mediates cellular proliferation and activation in renal myofibroblasts *in vitro* (related to Figure 4)**

(A) Representative western blot showing the levels of WWP2 isoforms in primary cultured renal myofibroblasts upon TGFβ1 stimulation (5ng/ml, 72h).

(B) Representative western blot showing the levels of WWP2 in primary cultured renal myofibroblasts that were transfected with WWP2 overexpression (WWP2$^{OE}$) or scrambled (Scram) DNAs.

(C) Representative western blot showing ECM protein levels in cultured WWP2$^{OE}$ and Scram myofibroblasts with or without TGFβ1 stimulation (5ng/ml, 72h).

(D*) Left:* representative graph of cell cycle in cultured WWP2$^{OE}$ and Scram myofibroblasts using flow cytometry (TGFβ1 stimulation (5ng/ml, 72h)). *Right*: quantification of cell cycle phases at G0/G1, S and G2/M in cultured WWP2$^{OE}$ and Scram myofibroblasts (n = 3, i.e., from 3 independent experiments). Values are reported as mean ± SEM.

### Supplementary Figure 6

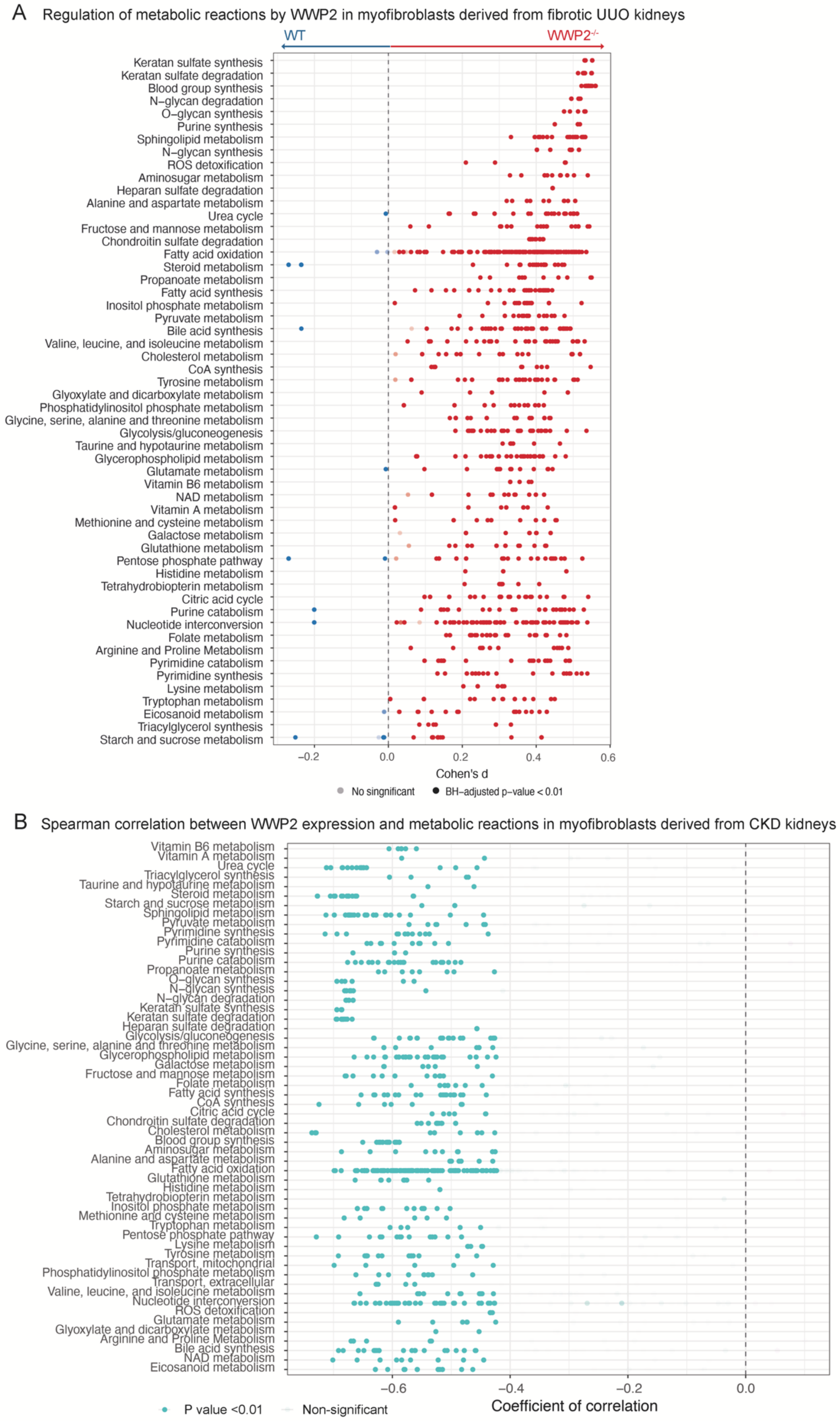

**Supplementary Figure 6. Compass-based exploration of metabolic profile (related to Figure 5).**

(A) Compass-score differential activity test from ECM-expressing myofibroblasts derived from WT and WWP2$^{-/-}$ UUO kidneys. Same analysis as shown in Figure 5A, but showing all metabolic reactions. BH-adjusted P-value, P-value corrected for multiple testing using the Benjamini-Hochberg method. For each metabolic reaction, the effect size of the change associated with WWP2 deficiency was estimated by Cohen's D (x-axis). The metabolic reactions upregulated in WWP2 deficient mouse kidney are indicated in red.

(B) Spearman correlation between Compass scores and the expression of WWP2 in renal myofibroblasts derived 5 CKD patients [1]. Same analysis as shown in Figure 5B, but showing all metabolic reactions. The nonsignificant correlations are shown in light grey. The metabolic reactions that are negatively correlated (P<0.01) with WWP2 expression in myofibroblasts are indicated in light blue.

### Supplementary Figure 7

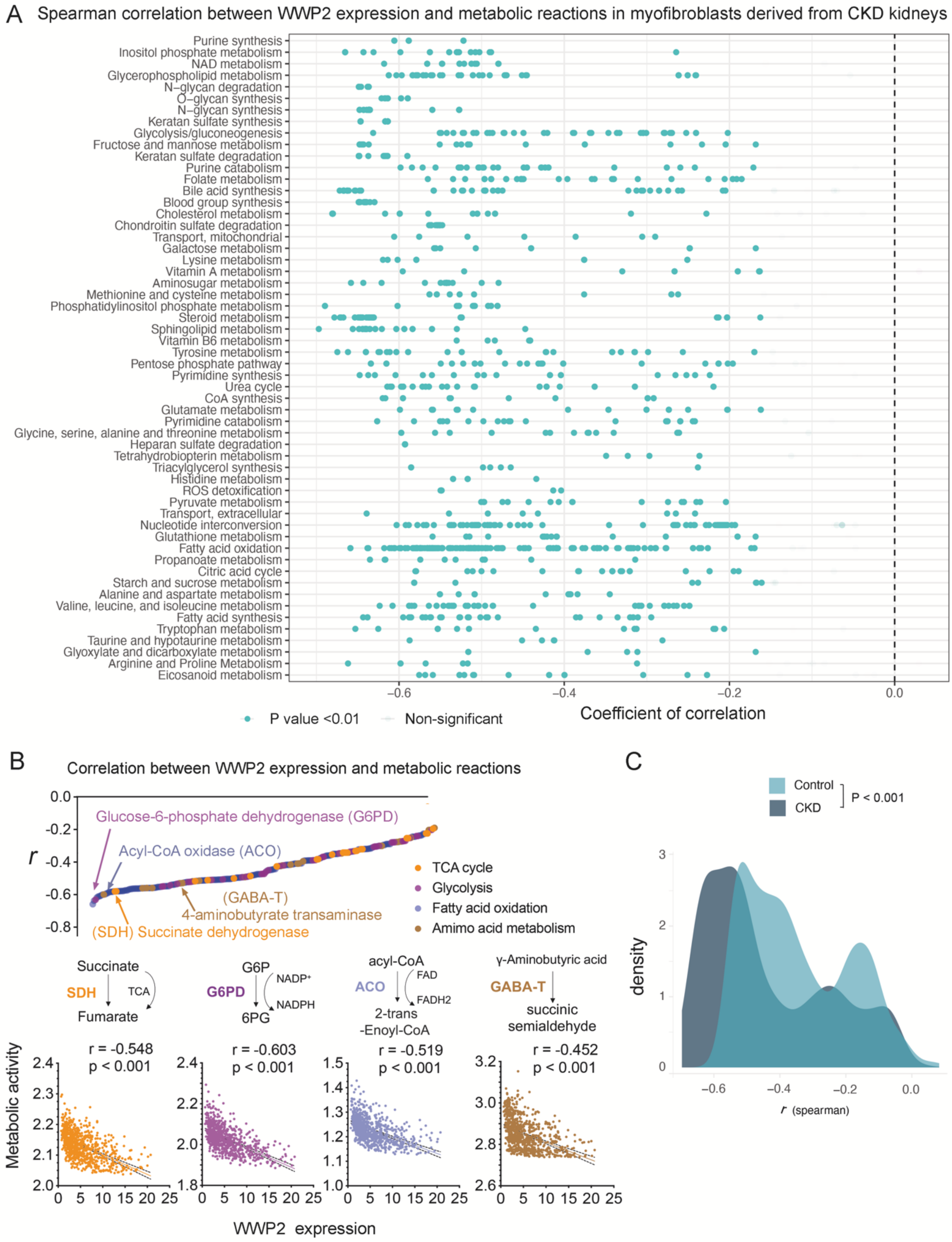


**Supplementary Figure 7. Compass-based exploration of metabolic reactions regulated by WWP2 in another CKD cohort (related to Figure 5).**

(A) Spearman correlation between Compass scores and the expression of WWP2 in renal myofibroblasts derived 10 patients with CKD [2]. Same analysis as shown in Supplementary Figure 6B (in 5 CKD patients) and nonsignificant correlations shown in light grey. The metabolic reactions that are negatively correlated ($P<0.01$) with WWP2 expression in myofibroblasts are indicated in light blue.

(B) Spearman correlation of Compass scores with WWP2 expression in renal myofibroblasts from 10 CKD patients. *Upper*: The significant Spearman correlation between Compass scores and the expression levels of WWP2 in renal myofibroblasts. The color coding represents different metabolic pathways. *Lower*: Spearman correlation analysis for 4 key metabolic reactions. The whole set of metabolic reactions was showed in supplementary Figure 7A.

(C) Histogram showing the distribution of Spearman correlations between WWP2 expression and metabolic actions in renal myofibroblasts derived from 12 controls and 10 CKD patients, showing that the correlation (association) is scientifically stronger ($P<0.001$) in CKD compared with control myofibroblasts. P-value was calculated using two-tailed Wilcoxon rank-sum test.

### Supplementary Figure 8

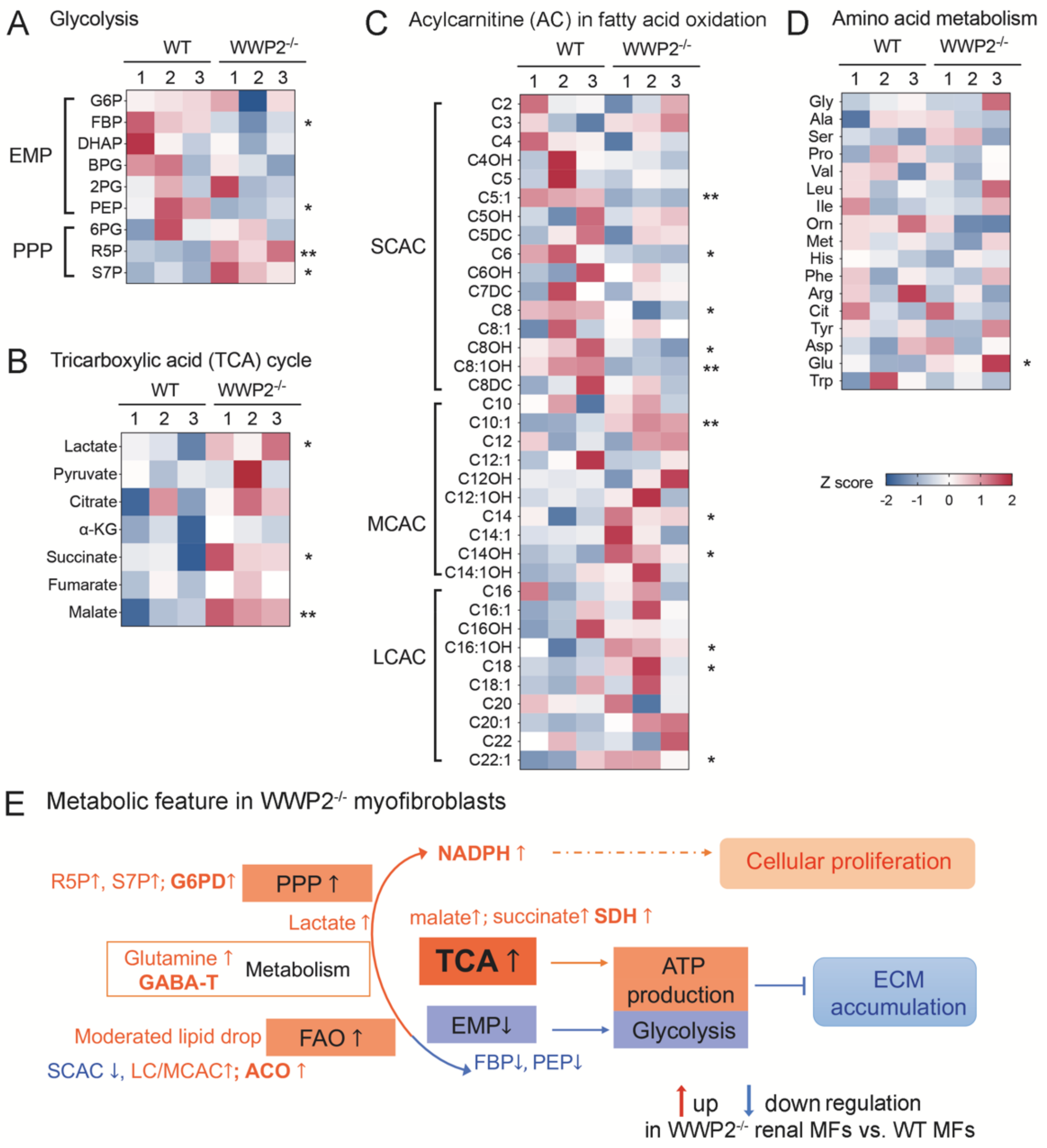


**Supplementary Figure 8. Profile of metabolites measured in cultured renal myofibroblasts derived from WT and WWP2$^{-/-}$ kidneys (related to Figure 5).**

(A-D) Metabolomics profile of metabolites in cultured renal myofibroblasts (P2) derived from WT and WWP2$^{-/-}$ kidneys, arranged according to metabolic pathways: glycolysis pathway (A), TCA cycle (B), ACs in fatty acid oxidation pathway (C), and amino acid metabolism (D). Same analysis as shown in Figure 5C, but showing all metabolites (and not just significantly differentially expressed between WT and WWP2$^{-/-}$, as in the main figure). Each metabolite was presented as Z score. (E). Schematic representation illustrating the metabolic processes changed in WWP2$^{-/-}$ myofibroblasts.

EMP: Embden-Meyerhof pathway of glycolysis; PPP: pentose phosphate pathway; FBP: Fructose-1,6-bisphosphate; PEP: Phosphoenolpyruvate; R5P: Ribose-5-phosphate; S7P: Sedoheptulose-7-phosphate; G6P: Glucose-6-phosphate; DHAP: Dihydroxyacetone phosphate; BPG: Bisphosphoglycerate or 2,3-Bisphosphoglycerate; 2PG: 2-Phosphoglycerate;

SCAC: small chain of acylcarnitine; LCAC: long chain of acylcarnitine; MCAC: Medium chain of acylcarnitine; α-KG, a-ketoglutarate; Gly: glycine; Ala: alanine; Ser: serine; Pro: proline; Val: valine; Leu: leucine; Ile: isoleucine; Orn: ornithine; Met: methionine; His: histidine; Phe: phenylalanine; Arg: arginine; Cit: citrate; Tyr: tyrosine; Asp: aspartic acid; Glu: glutamic acid; Trp: tryptophan.

### Supplementary Figure 9

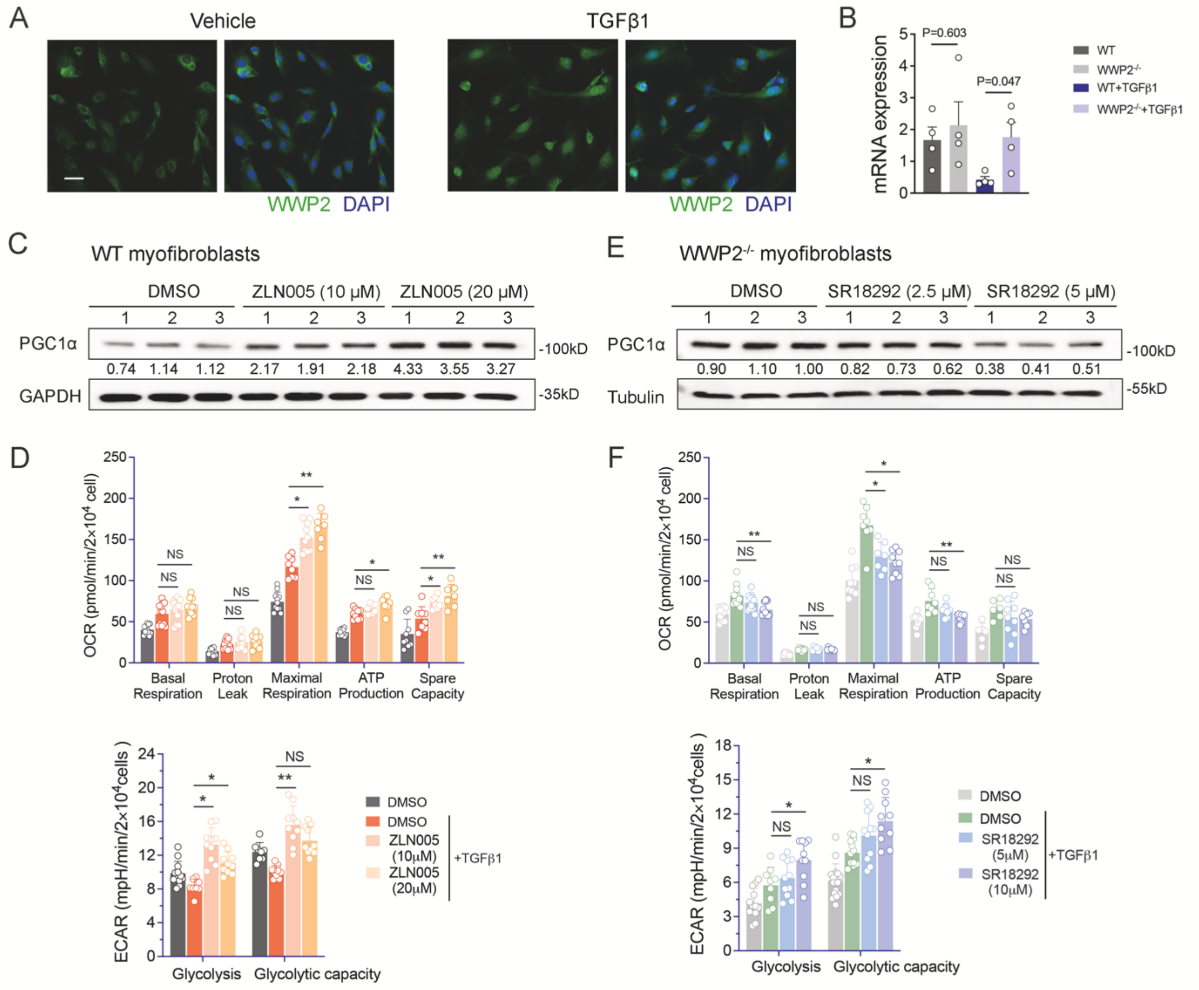


**Supplementary Figure 9. Regulation of PGC-1α signalling by WWP2 (related to Figure 6)**

(A) Representative immunofluorescence analysis of WWP2 in WT primary renal myofibroblasts showing nuclear localization after 24 hrs TGFβ1 stimulation (5ng/ml). Scale bar: 20 μm.

(B) Normalized gene expression levels (TPM, transcripts per million kilobase) measured by bulk-RNA seq analysis of PGC-1α in myofibroblasts derived from WT and WWP2$^{-/-}$ kidneys with or without TGFβ1 stimulation (5ng/ml, 72hrs). n=4, from independent biological replicates. Values are reported as mean ± SEM.

(C) Representative western blot showing the effect of ZLN005 (10 μM and 20 μM) on PGC-1α protein expression in cultured WT myofibroblasts following 24 hrs TGFβ1 (5ng/ml). DMSO (10 μM) was used as a reference control.

(D) Barplots summarizing the effects of ZLN005 (10 μM and 20 μM) on OCR based on Seahorse Mito stress assays (*upper panel*) and on ECAR based on Seahorse Glycolysis assays (*lower panel*) in cultured WT myofibroblasts. Three biological experiments were carried out, each consisting of independent cell plating on 3-4 Seahorse microplate wells (yielding n=9-12 data points).

(E) Representative western blot showing the effect of SR18292 (2.5 μM and 5 μM) on PGC-1α protein expression in cultured WWP2$^{-/-}$ myofibroblasts following 24 hrs TGFβ1 (5ng/ml). DMSO (5 μM) was used as a reference control.

(F) Barplots summarizing the effects of SR18292 (2.5 µM and 5 µM) on OCR based on Seahorse Mito stress assays (*upper panel*) and on ECAR based on Seahorse Glycolysis assays (*lower panel*) in cultured WWP2$^{-/-}$ myofibroblasts. Three biological experiments were carried out, each consisting of independent cell plating on 3-4 Seahorse microplate wells (yielding n=9-12 data points).
